# Clathrin-independent endocytic retrieval of SV proteins mediated by the clathrin adaptor AP-2 at mammalian central synapses

**DOI:** 10.1101/2021.06.24.449713

**Authors:** Tania López-Hernández, Koh-ichiro Takenaka, Yasunori Mori, Pornparn Kongpracha, Shushi Nagamori, Volker Haucke, Shigeo Takamori

## Abstract

Neurotransmission is based on the exocytic fusion of synaptic vesicles (SVs) followed by endocytic membrane retrieval and the reformation of SVs. Conflicting models have been proposed regarding the mechanisms of SV endocytosis, most notably clathrin/ AP-2-mediated endocytosis and clathrin-independent ultrafast endocytosis. Partitioning between these pathways has been suggested to be controlled by temperature and stimulus paradigm. We report on the comprehensive survey of six major SV proteins to show that SV endocytosis in hippocampal neurons at physiological temperature occurs independent of clathrin while the endocytic retrieval of a subset of SV proteins including the vesicular transporters for glutamate and GABA depend on sorting by the clathrin adaptor AP-2. Our findings highlight a clathrin-independent role of the clathrin adaptor AP-2 in the endocytic retrieval of select SV cargos from the presynaptic cell surface and suggest a unified model for the endocytosis of SV membranes at mammalian central synapses.

## INTRODUCTION

Synaptic transmission relies on the release of neurotransmitters by calcium-triggered exocytic fusion of synaptic vesicles (SVs), tiny organelles (∼ 40 nm in diameter) that store and secrete neurotransmitter molecules at specialized active zone (AZ) release sites within presynaptic nerve terminals (1). Following exocytosis, SVs are locally reformed via compensatory endocytic retrieval of membrane and its protein constituents (i.e SV proteins) from the plasma membrane in order to keep the presynaptic membrane surface area constant and to ensure sustained neurotransmission (2, 3). While the core components that mediate the exo- and endocytosis of SVs have been identified and characterized in detail (4–6), the molecular mechanisms underlying SV endocytosis and reformation have remained controversial (7–12).

Pioneering ultrastructural analyses of stimulated frog neuromuscular junctions using electron microscopy (EM) suggested that SVs are recycled via clathrin-mediated endocytosis (CME) of plasma membrane infoldings or budding from cisternal structures located away from the AZ (13). Subsequent studies in neurons and in non-neuronal models showed that CME occurs on a timescale of many seconds and crucially depends on clathrin and its essential adaptor protein complex 2 (AP-2)(14), a heterotetramer comprising α, β2, μ2, and σ2 subunits, as well as a plethora of endocytic accessory proteins (1-4, 15, 16). These and other works led to the view that SVs are primarily, if not exclusively, recycled by clathrin/ AP-2-dependent CME (17, 18).

Recent studies using high-pressure freezing EM paired with optogenetic stimulation have unravelled a clathrin-independent mechanism of SV endocytosis (CIE) in response to single action potential (AP) stimuli that selectively operates at physiological temperature (10). This ultrafast endocytosis (UFE) pathway is distinct from kiss-and-run or kiss-and-stay exo-endocytosis observed in neuroendocrine cells (19, 20), operates on a timescale of hundreds of milliseconds, and results in the generation of endosome-like vacuoles (ELVs) from which SVs can reform via clathrin-mediated budding processes (8, 11). Temperature-sensitive, clathrin-independent SV endocytosis has also been observed by presynaptic capacitance recordings at cerebellar mossy fiber boutons and by optical imaging at small hippocampal synapses (7–9) and is compatible with the accumulation of post-endocytic presynaptic vacuoles upon acute or sustained genetic perturbation of clathrin at stimulated fly neuromuscular junctions (21–23) and at mammalian central synapses (8, 24). Collectively, these studies suggest that SV endocytosis under physiological conditions is primarily mediated by CIE (e.g. UFE), while the function of clathrin and clathrin adaptors such as AP-2 is limited to the reformation of functional SVs from internal ELVs rather than acting at the plasma membrane proper.

While this model can provide a mechanistic explanation for the observed speed of SV endocytosis, the key question of how SV proteins are sorted to preserve the compositional integrity of SVs (5) remains unresolved. First, optical imaging-based acid-quench experiments in hippocampal neurons indicate that the capacity of UFE is limited to single or few APs, while the majority of SV proteins appear to be internalized on a timescale of several seconds following AP train stimulation (9); (2, 25), i.e. a timescale compatible with either CME or CIE. Second, as high-pressure freezing EM experiments have not been able to reveal the fate and time course of endocytosis of SV proteins, it is formally possible that UFE proceeds under conditions of clathrin depletion (11), while SV proteins remain stranded on the neuronal surface. Third, mutational inactivation of the binding motifs for the clathrin adaptor AP-2 in the vesicular transporters for glutamate (VGLUT) and γ-aminobutyric acid (VGAT) severely compromises the speed and efficacy of their endocytic retrieval at room temperature (26–29), arguing that at least under these conditions (e.g. low temperature when UFE is blocked) these SV proteins may be retrieved from the cell surface via clathrin/ AP-2. Finally, genetic inactivation of clathrin/ AP2-associated endocytic adaptors for the sorting of specific SV proteins, e.g. Stonin 2, an adaptor for the SV calcium sensor Synaptotagmin, or AP180, an adaptor for Synaptobrevin/ VAMPs, causes the accumulation of their respective SV cargos at the neuronal plasma membrane (3, 30–33). These data could be interpreted to indicate that at least some SV proteins are endocytosed via CME, whereas others may use CIE mechanisms such as UFE. However, such a model bears the problem of how CME and CIE pathways are coordinated and how membrane homeostasis is then maintained.

To solve the question how SV protein sorting is accomplished and how this relates to CME-*vs* CIE-based mechanisms for SV endocytosis, we have conducted a comprehensive survey of six major SV proteins in primary hippocampal neurons depleted of clathrin or conditionally lacking AP-2. We show that clathrin is dispensable for the endocytosis of all SV proteins at physiological temperature independent of the stimulation paradigm. In contrast, endocytic retrieval of a subset of SV proteins including VGLUT1 and VGAT depends on sorting by AP-2. Our findings highlight a clathrin-independent function of the clathrin adaptor AP-2 in the endocytic retrieval of select SV cargos from the presynaptic plasma membrane and suggest a unified model for SV endocytosis and recycling.

## RESULTS

Based on prior works (2, 4, 7–9, 11–13, 15, 34) three main models for the sorting and endocytic recycling of SV proteins at central mammalian synapses can be envisaged (Figure 1). According to the classical CME-based model of SV endocytosis, SV proteins exocytosed in response to AP trains undergo clathrin/ AP-2-mediated sorting and endocytosis from the presynaptic plasma membrane or plasma membrane infoldings (34) akin to CME in receptor-mediated endocytosis in non-neuronal cells (16). This model predicts that loss of either clathrin or its essential adaptor protein complex AP-2 delays the endocytic retrieval of all major SV proteins (Figure 1A). A second model supported by elegant high-pressure freezing (11), electrophysiological (7), and optical imaging (8, 9) experiments suggests that exocytosed SV proteins are internalized via clathrin- and AP-2-independent bulk endocytosis. In this model, SV protein sorting occurs from internal ELVs that are formed downstream of the endocytic internalization step. Hence, at physiological temperature the endocytic retrieval of all major SV proteins would proceed unperturbed in the absence of either clathrin or AP-2 (Figure 1B). Finally, it is conceivable that exocytosed SV proteins present on the neuronal surface are sorted by dedicated endocytic adaptors, e.g. the AP-2 complex, to facilitate their clathrin-independent internalization via CIE. Clathrin, possibly in conjunction with AP-2 and other adaptors then operates downstream of CIE to reform functional SVs from ELVs. In this case, loss of clathrin or AP-2 are predicted to result in distinct phenotypes: While endocytosis of SV proteins is unperturbed upon depletion of clathrin, loss of AP-2 would be expected to selectively affect the rate and efficacy of endocytosis of distinct SV cargos recognized by AP-2 (Figure 1C).

**Figure 1.**
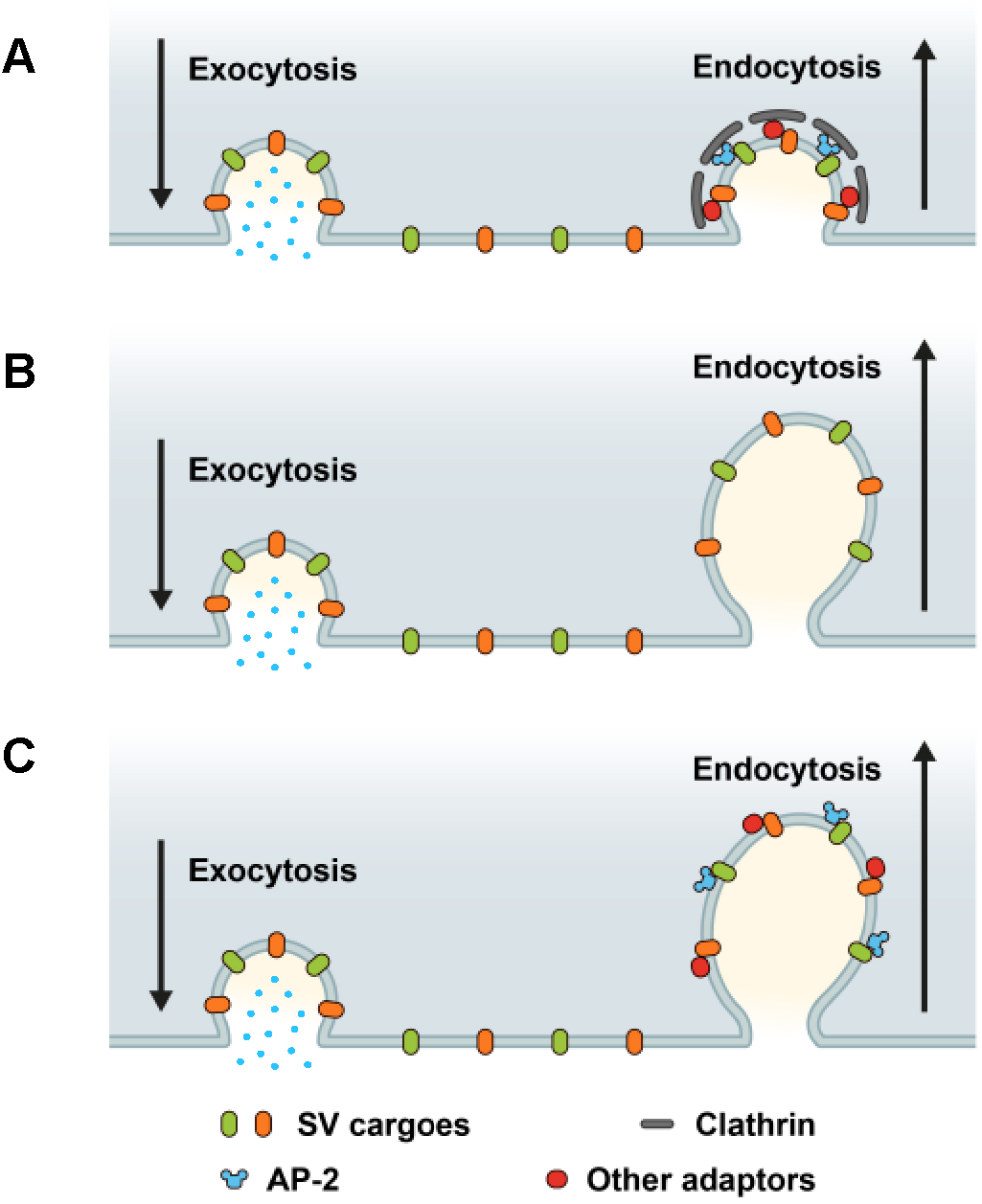
Possible roles of clathrin and AP-2 in SV endocytosis and SV cargo retrieval. (A) A model predicting that SV retrieval following neurotransmitter release is mediated by clathrin mediated endocytosis (CME) where AP-2 functions as bridge between SV cargos and clathrin to form clathrin-coated pits (CCP) at plasma membrane. In this scenario, inactivation of both clathrin and AP-2 would slow either SV endocytosis as well as SV cargo retrieval. (B) A model predicting that SV endocytosis occurs in a clathrin-independent manner (CIE), and neither clathrin nor AP-2 mediate SV endocytosis and SV cargo retrieval at plasma membrane. If this were the case, inactivation of both clathrin and AP-2 would not change the kinetics rate of SV endocytosis and SV cargo retrieval. (C) A model predicting that dedicated adaptors such as AP-2 function as sorting protein for SV cargo even during CIE. If this were the case, inactivation of clathrin and AP-2 would produce distinct phenotypes between SV endocytosis and SV cargo retrieval.

### Endocytic retrieval of SV proteins in hippocampal neurons occurs independent of clathrin at physiological temperature

To distinguish between these models, we optically recorded the stimulation-induced exo-endocytosis of SV proteins carrying within their lumenal domains a pH-sensitive super-ecliptic green fluorescent protein (SEP, often also referred to as pHluorin) that is de-quenched during exocytosis and undergoes re-quenching as SVs are internalized and re-acidified (35, 36). Specifically, we monitored SEP-tagged chimeras of the calcium sensor Synaptotagmin 1 (Syt1), the multi-spanning glycoprotein SV2A, the SNARE protein VAMP/Synaptobrevin 2 (hereafter referred to as Syb2), the tetraspanin Synaptophysin (Syp), the vesicular glutamate transporter 1 (VGLUT1), and the vesicular GABA transporter (VGAT), which have been used extensively to monitor SV recycling in various preparations (8, 9, 17, 25, 26, 35, 36) and constitute the major protein complement of SVs based on their copy numbers (5).We capitalized on the fact that in hippocampal neurons stimulated with trains of APs SV endocytosis occurs on a timescale of >10 seconds at physiological temperature (9), e.g. a time scale that is much slower than re-quenching of SEP due to reacidification of newly endocytosed vesicles (37, 38). Therefore, under these conditions, the decay of SEP signals can serve as a measure of the time course of SV endocytosis.

We first depleted clathrin heavy chain (CHC) in hippocampal neurons using lentiviral vectors to ∼ 25% of the levels found in controls as evidenced by confocal imaging of immunostained samples (Figure 2-figure supplement 1A,B) and in agreement with previous data (8, 11, 17). To assess the effects of clathrin loss on the stimulation-induced endocytic retrieval of SV proteins, we stimulated control or clathrin-depleted hippocampal neurons expressing any one of the six major SEP-tagged SV proteins with a high-frequency stimulus train (200 APs applied at 40 Hz) at physiological temperature (35 ± 2°C) (hereafter abbreviated as PT), and monitored fluorescence rise and decay over time. Strikingly, exo-/endocytosis of all SEP-tagged SV proteins proceeded with unaltered kinetics, i.e. τ∼15–20 s, irrespective of the depletion of clathrin (Figure 2A-L). Similar results were seen if clathrin function was acutely blocked by application of the small molecule inhibitor Pitstop 2 (39) (Figure 2-figure supplement 1C,D). If these experiments were repeated under conditions of low-frequency stimulation (200 APs applied at 5 Hz) at room temperature (RT), i.e. conditions in which the efficacy of CIE is reduced (11), the endocytic retrieval of Syp-SEP (also often referred to as SypHy) or VGLUT1-SEP was delayed in neurons depleted of clathrin (Figure 2-figure supplement 1E-H), consistent with earlier data using Syt1-SEP as a reporter (8).

**Figure 2.**
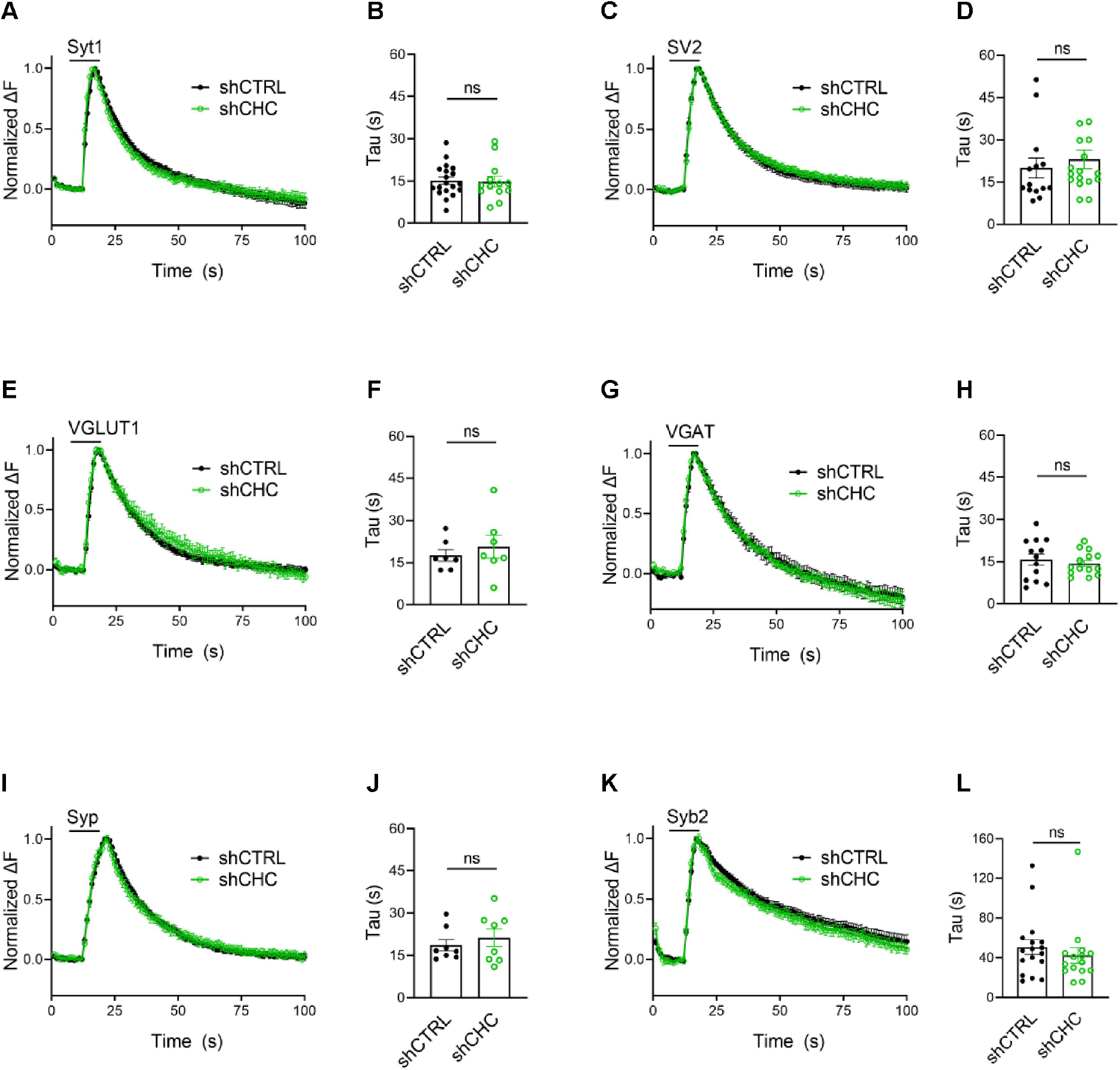
SV endocytosis in hippocampal neurons occurs independent of clathrin at PT. Average normalized traces of neurons transduced with lentivirus expressing non-specific shRNA (shCTRL) or shRNA targeting CHC (shCHC) and co-transfected with SEP probes tagged to the luminal portion of Syt1 (A), SV2 (C), VGLUT1 (E), VGAT (G), Syp (I) and Syb2 (K) subjected to electrical stimulation of 40 Hz (200 APs) at PT. Endocytic decay constant (Tau) of transfected and lentivirally transduced neurons co-expressing respectively (B) Syt1-SEP and shCTRL (15.2 ± 1.30 s) or shCHC (14.9 ± 1.87 s); (D) SV2-SEP and shCTRL (20.0 ± 3.53 s) or shCHC (23.1 ± 3.31 s); (F) VGLUT1-SEP and shCTRL (17.7 ± 2.05 s) or shCHC (20.8 ± 4.07 s); (H) VGAT-SEP and and shCTRL (15.8 ± 2.04 s) or shCHC (14.4 ± 1.14 s); (J) Syp-SEP and shCTRL (18.7 ± 2.03 s) or shCHC (21.3 ± 3.10 s); and (L) Syb2-SEP and shCTRL (50.6 ± 7.49 s) or shCHC (42.3 ± 8.03 s). Data shown represent the mean ± SEM for Syt1 (n_CTRL_ = 19 images, n_shCHC_ = 13 images; *p* = 0.875s), for SV2 (n_shCTRL_ = 14 images, n_shCHC_ = 17 images; *p* = 0.533), for VGLUT1 (n_shCTRL_ = 7 images, n_shCHC_ = 7 images; *p* = 0.506), for VGAT (n_shCTRL_ = 13 images, n_shCHC_ = 14 images; *p* = 0.534), for Syp (n_shCTRL_ = 8 images, n_shCHC_ = 8 images; *p* = 0.490) and for Syb2 (n_shCTRL_ = 17 images, n_shCHC_ = 15 images; *p* = 0.455). Two-sided unpaired *t*-test. Raw data can be found in Figure 2-source data 1.

These results show that at small hippocampal synapses at PT, endocytosis of all major SV proteins and hence, of SVs as a whole, occurs independent of clathrin via CIE.

### Clathrin-independent endocytosis of a subset of SV proteins depends on the clathrin adaptor AP-2

We next set out to analyze whether endocytosis of the major SV proteins is also independent of the essential clathrin adaptor complex AP-2 as expected, if SV endocytosis was mediated by CIE and the sole function of clathrin/ AP-2 was to reform SVs from postendocytic ELVs (Figure 1B). We conditionally ablated AP-2 expression by tamoxifen induction of Cre recombinase in hippocampal neurons from AP-2μ^lox/lox^ mice resulting in a reduction of AP-2 levels to < 15% of that detected in WT control neurons (hereafter referred to as AP-2µ KO) (Figure 3-figure supplement 2A,B) (8, 9). Further depletion below this level caused neuronal death.

Endocytosis of Synaptotagmin 1-SEP and SV2A-SEP proceeded with similar kinetics in control or AP-2μ KO hippocampal neurons stimulated with 200 APs applied at 40 Hz at physiological temperature, consistent with our earlier findings (8, 9) (Figure 3A-D). Surprisingly, however, we found that loss of AP-2 significantly slowed down the endocytic retrieval of other major SV proteins such as Synaptophysin, Synaptobrevin 2, and most prominently, of the vesicular neurotransmitter transporters VGLUT1 and VGAT (Figure 3E-L). These phenotypes were specific as plasmid-based re-expression of AP-2μ in AP-2μ KO neurons rescued defective endocytosis of these SEP-tagged SV proteins (Figure 3).

**Figure 3.**
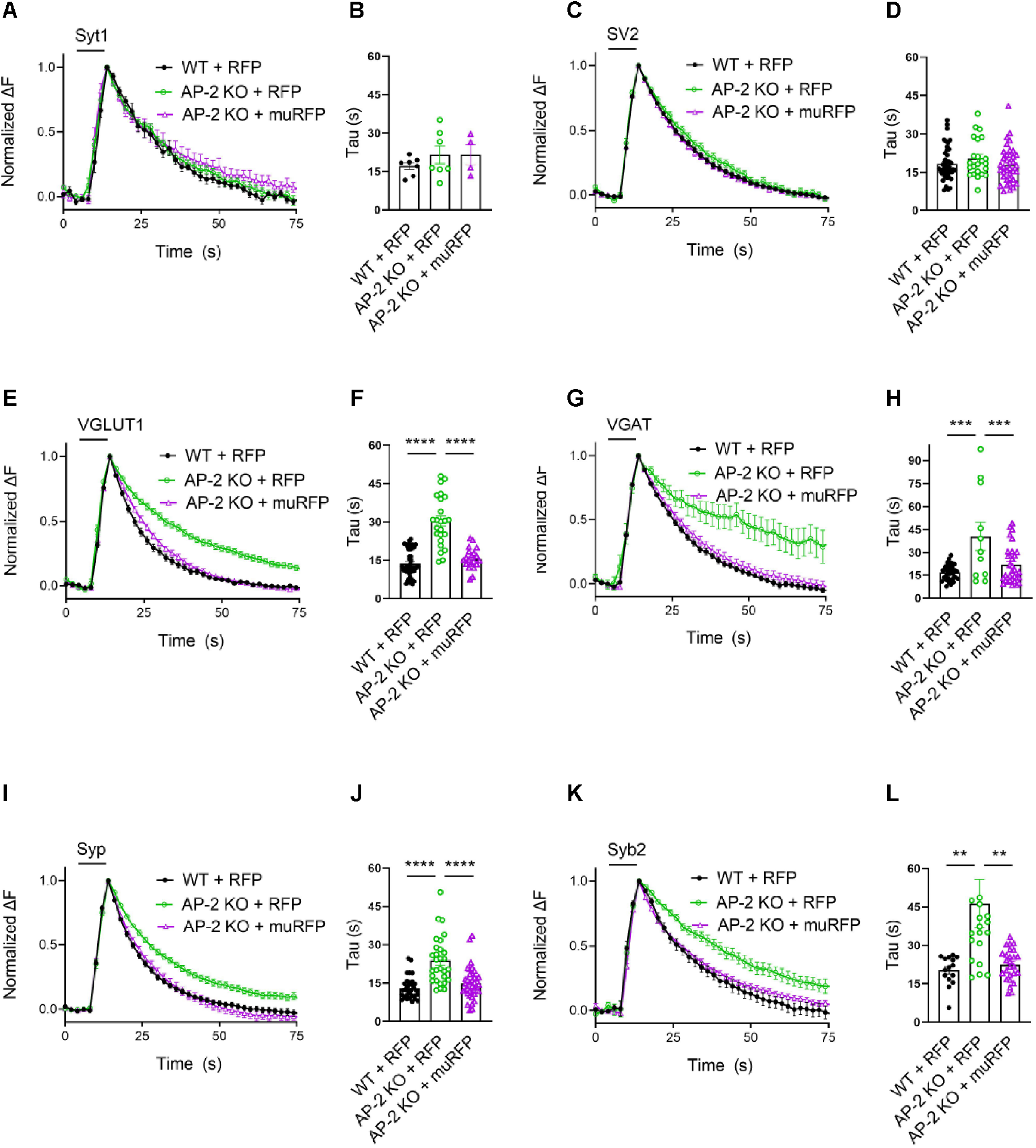
Clathrin-independent endocytic retrieval of select SV cargos by the clathrin adaptor AP-2 at PT. (A-D) Post-stimulus membrane retrieval of Syt1 and SV2 in the absence of AP-2 persists unaffected in response to 200 AP applied at 40 Hz. Average normalized traces of WT and AP-2 KO derived neurons co-transfected with Syt1-SEP (A) or SV2-SEP (C) and RFP or rescued by re-expression of untagged AP-2µ subunit together with soluble RFP (muRFP) in response to 200 AP applied at 40 Hz. Quantification of the endocytic decay constant (Tau) of neurons expressing Syt1-SEP (B) (τ_WT+RFP_ = 17.17 ± 1.347 s, τ_AP2 KO+RFP_ = 21.53 ± 3.430 s, τ_AP2 KO+mu-RFP_ = 21.51 ± 3.908 s) or SV2-SEP (D) (τ_WT+RFP_ =18.33 ± 0.988 s, τ_AP2 KO+RFP_ = 20.45 ± 1.56 s, τ_AP2 KO+mu-RFP_ = 18.04 ± 1.16 s). Data shown represent the mean ± SEM: Syt1 (n_WT+RFP_ = 7 images, n_AP2 KO+RFP_ = 7 images, n_AP2 KO+mu-RFP_ = 4 images; *p*_(WT+RFP vs AP2KO+RFP)_ = 0.4994; *p*_(AP2KO+RFP vs AP2KO+muRFP) >_ 0.9999); SV2 (n_WT+RFP_ = 44 images, n_AP2 KO+RFP_ = 23 images, n_AP2 KO+mu-RFP_ = 37 images; *p*_(WT+RFP vs AP2KO+RFP)_ = 0.4665; *p*_(AP2KO+RFP vs AP2 KO+mu-RFP)_=0.3943). One-way ANOVA with Tukey’s post-test. (E-L) Loss of AP-2 significantly delay the endocytic retrieval of other major SV proteins. Average normalized traces of WT and AP-2 KO neurons co-transfected with VGLUT1-SEP (E), VGAT-SEP (G), Syp-SEP (I), Syb2-SEP (K) and RFP or muRFP to rescue AP-2μ expression stimulated with 200 AP at 40 Hz. Time constants (Tau) were calculated from WT and AP-2 KO neurons expressing VGLUT1-SEP (F) (τ_WT+RFP_ = 13.52 ± 0.903 s, τ_AP2 KO+RFP_ = 30.33 ± 2.011 s, τ_AP2 KO+mu-RFP_ = 16.38 ± 1.002 s), VGAT-SEP (H) (τ_WT+RFP_ =16.76 ± 0.869 s, τ_AP2 KO+RFP_ = 40.56 ± 9.211 s, τ_AP2 KO+mu-RFP_ = 21.77 ± 2.148 s), Syp-SEP (J) (τ_WT+RFP_ =13.05 ± 0.743 s, τ_AP2 KO+RFP_ = 23.76 ± 1.719 s, τ_AP2 KO+mu-RFP_ = 14.89 ± 1.035 s), VGAT-SEP (D) (τ_WT+RFP_ =18.33 ± 0.988 s, τ_AP2 KO+RFP_ = 20.45 ± 1.56 s, τ_AP2 KO+mu-RFP_ = 18.04 ± 1.16 s), Syb2-SEP (L) (τ_WT+RFP_ =20.27 ± 1.476 s, τ_AP2 KO+RFP_ = 46.34 ± 9.425 s, τ_AP2 KO+mu-RFP_= 22.46 ± 1.211 s). Data shown represent the mean ± SEM: VGLUT1 (n_WT+RFP_ = 37 images, n_AP2 KO+RFP_ = 24 images, n_AP2 KO+mu-RFP_ = 23 images; \*\*\*\**p*_(WT+RFP vs AP2 KO+RFP)_ < 0.0001; \*\*\*\**p*_(AP2 KO+RFP vs AP2 KO+mu-RFP)_ < 0.0001); VGAT (n_WT+RFP_ = 34 images, n_AP2 KO+RFP_ = 11 images, n_AP2 KO+mu-RFP_ = 32 images; \*\*\*\**p*_(WT+RFP vs AP2 KO+RFP)_ < 0.0001; \*\*\**p*_(AP2 KO+RFP vs AP2 KO+mu-RFP)_ = 0.0008); Syp (n_WT+RFP_ = 33 images, n_AP2 KO+RFP_ = 29 images, n_AP2 KO+mu-RFP_ = 37 images; \*\*\*\**p*_(WT+RFP vs AP2 KO+RFP)_ < 0.0001; \*\*\*\**p*_(AP2 KO+RFP vs AP2 KO+mu-RFP)_ < 0.0001); Syb2 (n_WT+RFP_ = 15 images, n_AP2 KO+RFP_ = 20 images, n_AP2 KO+mu-RFP_ = 26 images; \*\**p*_(WT+RFP vs AP2 KO+RFP)_ = 0.0083; \*\**p*_(AP 2KO+RFP vs AP2 KO+mu-RFP)_ = 0.0052). One-way ANOVA with Tukey’s post-test. Raw data can be found in Figure 3-source data 1.

We challenged these data by monitoring SV endocytosis in response to a milder stimulation paradigm that results in the exocytic fusion of the readily-releasable pool of SVs (40, 41), i.e. 50 APs applied at 20 Hz. While endocytosis of SV2A- and VGLUT1-SEP proceeded unperturbed in hippocampal neurons depleted of clathrin (Figure 4A-D), a substantial delay in the endocytosis of VGLUT1- but not SV2-SEP was observed in neurons lacking AP-2 (Figure 4E-H). No difference was found in the fraction of boutons responding to stimulation with 50 APs between WT and AP-2μ KO neurons (Figure 4-figure supplement 3A,B). At lower stimulation intensities (i.e. 10 or 20 APs), AP-2μ KO neurons displayed significantly attenuated exocytic responses (Figure 4-figure supplement 3B), possibly reflecting a reduced release probability originating from defects in SV reformation, akin to the reported phenotype of clathrin loss in hippocampal neurons (10, 11).

**Figure 4.**
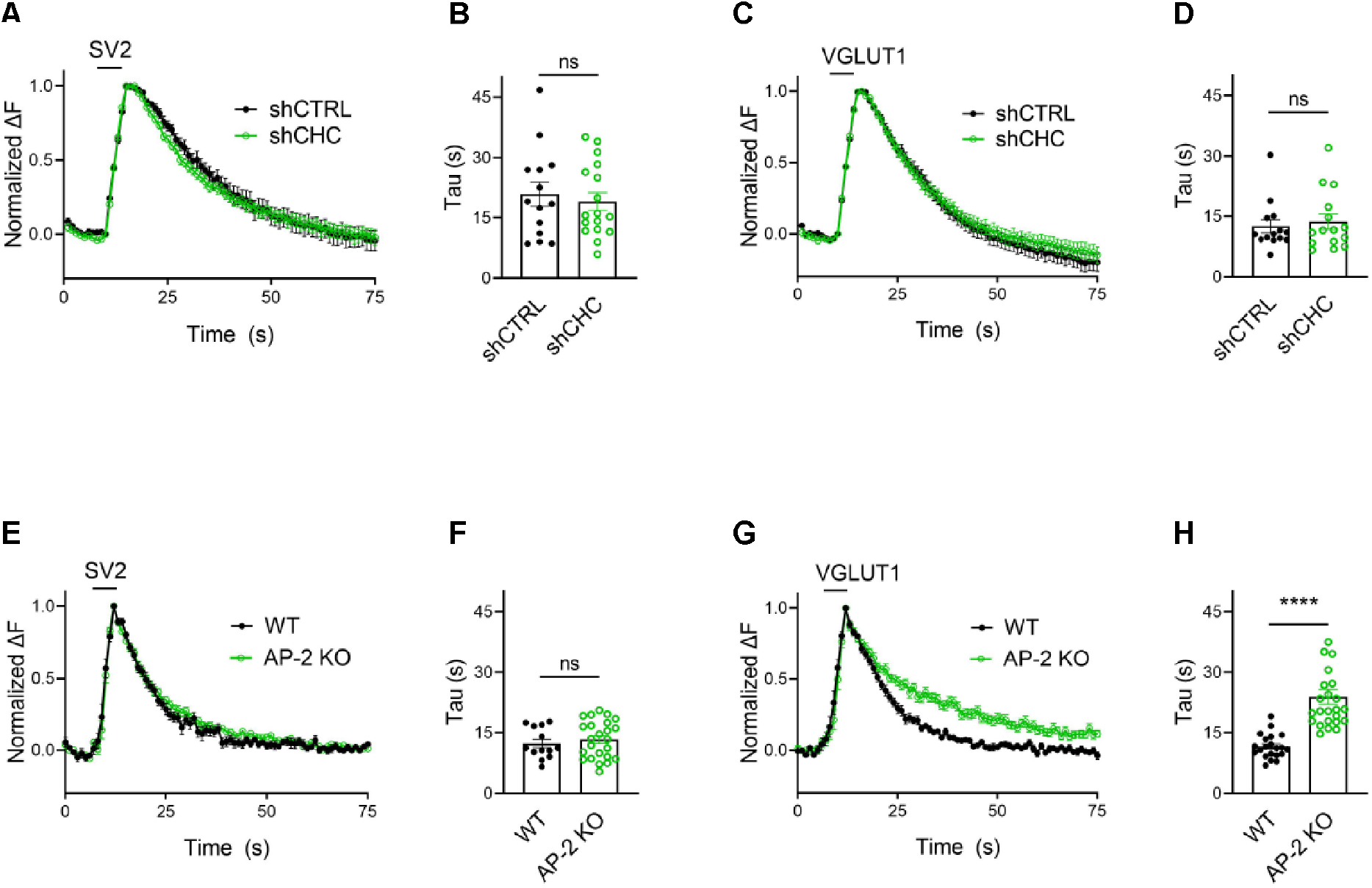
Clathrin-independent endocytic retrieval of SV proteins mediated by AP-2 is independent of the stimulation strength at PT. (A-D) Lack of clathrin does not alter the endocytosis of SV2 and VGLUT1 in response to stimulation with 50 APs (i.e. a stimulus that releases the RRP) at PT. Average normalized traces of neurons transduced with lentivirus expressing non-specific shRNA (shCTRL) or shRNA targeting CHC (shCHC) and co-transfected with either SEP-tagged SV2 (A) or VGLUT1 (C) stimulated with 50 AP applied at 20 Hz at PT. Quantification of the endocytic decay constant (Tau) in neurons co-expressing SV2 (B) and shCTRL (20.97± 2.991 s) or shCHC (19.18 ± 2.216 s); and VGLUT1 (D) and shCTRL (12.51 ± 1.621 s) or shCHC (13.68 ± 1.897 s). Data represent the mean ± SEM for SV2 (n_shCTRL_ = 14 images, n_shCHC_ = 17 images; *p* = 0.6274) and for VGLUT1 (n_CTRL_ = 14 images, n_CHC-KD_ = 15 images; *p* = 0.6468). Two-sided unpaired *t*-test. (E-H) Endocytosis delay for VGLUT1 but not for SV2 in neurons depleted of AP-2 when stimulated with a mild train of 50 AP. Average normalized traces of neurons from WT and AP-2 KO mice transfected with either SV2-SEP (E) or VGLUT1-SEP (G) in response of 50 AP applied at 20 Hz at PT. Quantification of the endocytic decay constant (Tau) of SV2-SEP-expressing neurons (F) (τ_WT_ = 12.40 ± 1.050 s, τ_AP2 KO_ = 13.34 ± 0.946 s) or VGLUT1-SEP (F) (τ_WT_ = 11.61 ± 0.651 s, τ_AP2 KO_ = 23.96 ± 1.871 s). Data represent the mean ± SEM for SV2 (n_WT_ = 13 images, n_AP2 KO_ = 24 images; *p* = 0.5355) and for VGLUT1 (n_WT_ = 21 images, n_AP2 KO_ = 24 images; \*\*\*\**p* < 0.0001). Two-sided unpaired *t*-test. Raw data can be found in Figure 4-source data 1.

These data unravel a clathrin-independent role of the clathrin adaptor AP-2 in the endocytic retrieval of select SV cargos including VGLUT1 and VGAT at physiological temperature, while endocytosis of Syt1 or SV2A proceeds with unaltered kinetics in the absence of AP-2.

### Clathrin-independent endocytosis of endogenous VGAT depends on the clathrin adaptor AP-2

As optical imaging of SEP reporters may lead to artifacts caused by overexpression of exogenous SV proteins (42), we analyzed the internalization kinetics of endogenous VGAT using antibodies directed against its luminal domain coupled to the pH-sensitive fluorophore cypHer 5E. The cyanine-based dye cypHer 5E is quenched at neutral pH but exhibits bright fluorescence when present in the acidic lumen of SVs (40) and, thus can serve as a tracer for the endocytosis of endogenous SV proteins when it is internalized into SVs prior to measurements (Figure 5A). First, we probed the effects of AP-2μ KO on VGAT endocytosis. Loss of AP-2 severely delayed the endocytic retrieval of endogenous VGAT in response to train stimulation with either 200 AP (Figure 5B,C) or 50 APs (Figure 5F,G) at physiological temperature, consistent with our results from exogenously expressed VGAT-SEP (see Figure 3).To determine whether the requirement for AP-2 reflects a function for CME in the retrieval of endogenous VGAT, we examined the effects of genetic or pharmacological blockade of clathrin function. Lentiviral shRNA-mediated depletion of clathrin (Figure 5-figure supplement 4A-D) potently blocked CME of transferrin (Figure 5-figure supplement 4E,F) but had no effect on the endocytic retrieval of endogenous VGAT in response to either strong (e.g. train of 200 AP applied at 40 Hz) (Figure 5D,E) or mild stimulation (50 AP at 20 Hz) (Figure 5H,I) at physiological temperature. Similar results were obtained, if clathrin function was perturbed pharmacologically by acute inhibition in the presence of Pitstop 2 (Figure 5B,C,F,G; Figure 5-figure supplement 4G,H). However, clathrin loss resulted in a significant reduction of the readily-releasable and total recycling vesicle pool sizes probed by consecutive trains of 50 APs and 900 APs interspersed by a 90 s inter-stimulus interval (6) (Figure 5-figure supplement 4I). In contrast, the kinetics of VGAT endocytosis was unaffected by clathrin loss under these conditions (Figure 5-figure supplement 4J).

**Figure 5.**
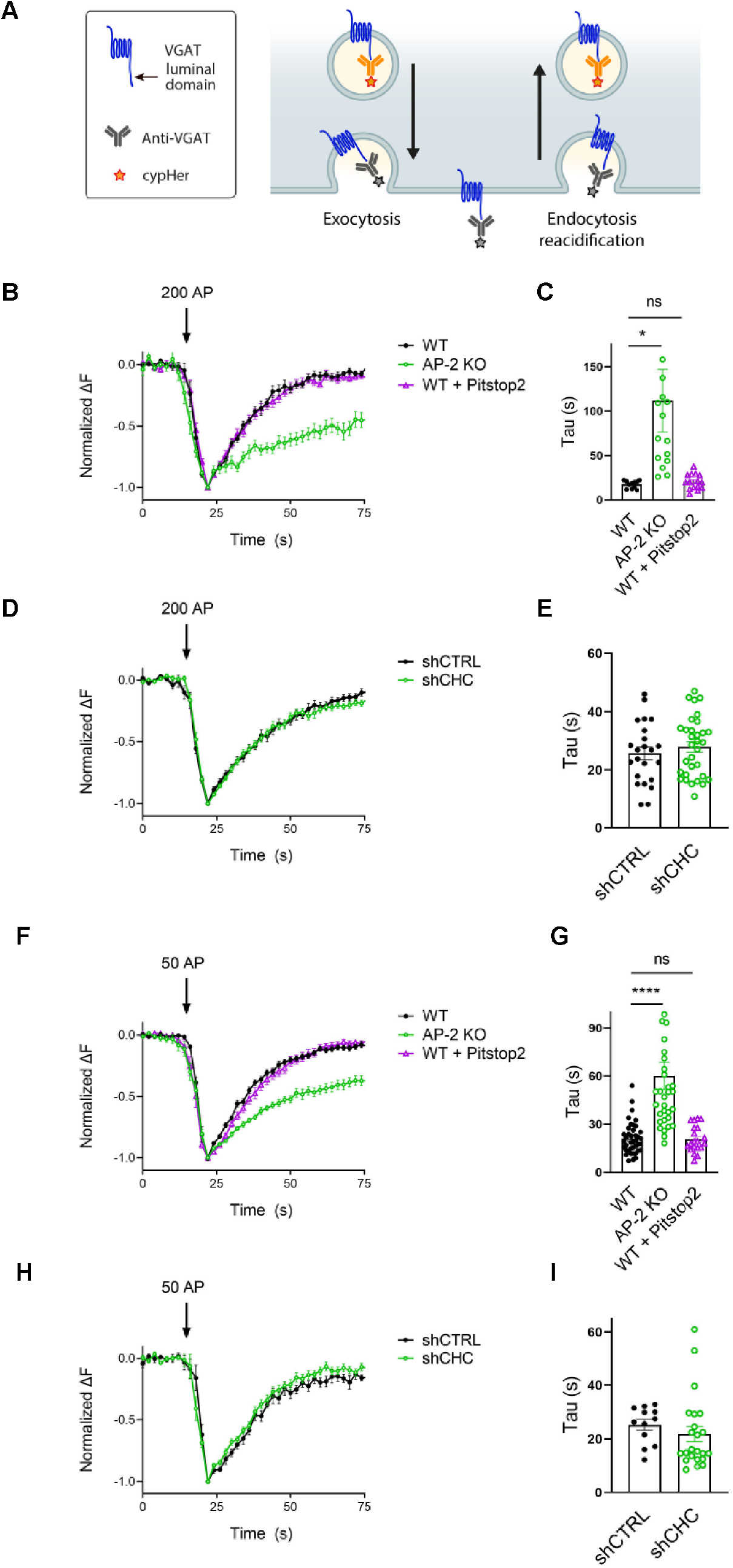
Post-exocytic sorting of endogenous VGAT depends on AP-2 but not clathrin at PT. (A) Diagram depicting the use of cypHer-coupled antibodies targeting the luminal domain of VGAT to monitor fluorescence changes during exo-endocytosis of endogenously labelled VGAT, as cypHer is a pH-sensitive fluorophore which is quenched at neutral extracellular pH. (B-I) Clathrin but not AP-2 is dispensable for endocytic retrieval of endogenous cypHer-labelled VGAT independent of the stimulation intensity at PT. (B) Average normalized traces of neurons from WT treated or not with the clathrin inhibitor Pitstop 2 and from AP-2 KO mice incubated with anti-VGAT cypHer-coupled antibodies for live labeling of synapses in response to a high-frequency stimulus train (200 AP at 40 Hz) at PT. (C) Quantification of the endocytic decay constant (Tau) of anti-VGAT cypHer-labelled neurons (τ_WT_ = 17.38 ± 1.303 s, τ_AP2 KO_ = 111.7 ± 35.12 s, τ_WT+Pistop2_ = 20.39 ± 2.238). Data represent the mean ± SEM for n_WT_ = 11 images, n_AP2 KO_ = 15 images and n_WT+Pistop2_ = 15 images. \**p*_WT vs AP2 KO_ = 0.0179, *p*_WT vs WT+Pitstop2_ = 0.9954, one-way ANOVA with Tukey’s post-test. (D) Average normalized traces of neurons transduced with lentivirus expressing non-specific shRNA (shCTRL) or shRNA targeting CHC (shCHC) incubated with anti-VGAT cypHer-coupled antibodies in response to a high-frequency stimulus train (200 AP at 40 Hz) at PT. (E) Quantification of the endocytic decay constant (Tau) of anti-VGAT cypHer-labelled neurons (τ_shCTRL_ = 25.71 ± 2.190 s, τ_shCHC_ = 27.90 ± 1.793 s). Data represent the mean ± SEM for n_shCTRL_ = 23 images and n_shCHC_= 32 images. *p* = 0.4395, two-sided unpaired *t*-test. (F) Average normalized traces of neurons from WT treated or not with the clathrin inhibitor Pitstop 2 and from AP-2 KO mice incubated with anti-VGAT cypHer-coupled antibodies for live labeling of synapses in response to a mild-frequency stimulus train (50 AP at 20 Hz) at PT. (G) Quantification of the endocytic decay constant (Tau) of anti-VGAT cypHer-labelled neurons (τ_WT_ = 21.13 ± 1.565 s, τ_AP2 KO_ = 60.04 ± 8.387 s, τ_WT+Pistop2_ = 20.73 ± 1.812). Data represent the mean ± SEM for n_WT_ = 40 images, n_AP2 KO_ = 32 images and n_WT+Pistop2_ = 20 images. \*\*\*\**p*_WT vs AP2 KO_ < 0.0001, *p*_WT vs WT+Pitstop2_ = 0.9986, one-way ANOVA with Tukey’s post-test. (H) Average normalized traces of neurons transduced with lentivirus expressing non-specific shRNA (shCTRL) or shRNA targeting CHC (shCHC) incubated with anti-VGAT cypHer-coupled antibodies in response to a mild-frequency stimulus train (50 AP at 20 Hz) at PT. (I) Quantification of the endocytic decay constant (Tau) of anti-VGAT cypHer-labelled neurons (τ_shCTRL_ = 25.23 ± 1.994 s, τ_shCHC_ = 21.84 ± 2.836 s). Data represent the mean ± SEM for n_shCTRL_ = 12 images and n_shCHC_ = 23 images. *p* = 0.4267, two-sided unpaired *t*-test. Raw data can be found in Figure 5-source data 1.

We conclude that at physiological temperature the endocytosis of endogenous VGAT from the neuronal surface depends on the clathrin adaptor AP-2 while clathrin function is dispensable. Instead, clathrin may facilitate the reformation of functional SVs from ELVs downstream of CIE to sustain neurotransmission (8, 11).

### AP-2 depletion causes surface stranding of endogenous vesicular neurotransmitter transporters but not of Syt1 and SV2A

As endocytosis of a subset of SV proteins, e.g. VGLUT1 and VGAT, was impaired in the absence of AP-2, one might expect their partial redistribution to the neuronal surface in AP-2μ KO neurons. To test this, we labeled surface-stranded SV proteins by a membrane-impermeant biotinylating reagent in cultured cerebellar granule neurons derived from AP-2μ KO mice or wild-type littermate controls. Biotinylated proteins were captured on a streptavidin matrix and analyzed by immunoblotting (Figure 6A). No difference was detected in the plasma membrane levels of Syt1 and SV2A between WT and AP-2μ KO neurons (Figure 6B-E). By contrast, significantly larger amounts of VGLUT1 and VGAT were found at the plasma membrane of AP-2μ KO neurons compared to WT controls (Figure 6F-I), while the total levels of SV proteins assessed either by western blot or immunostaining were unaltered (Figure 6-figure supplement 5A,B).

**Figure 6.**
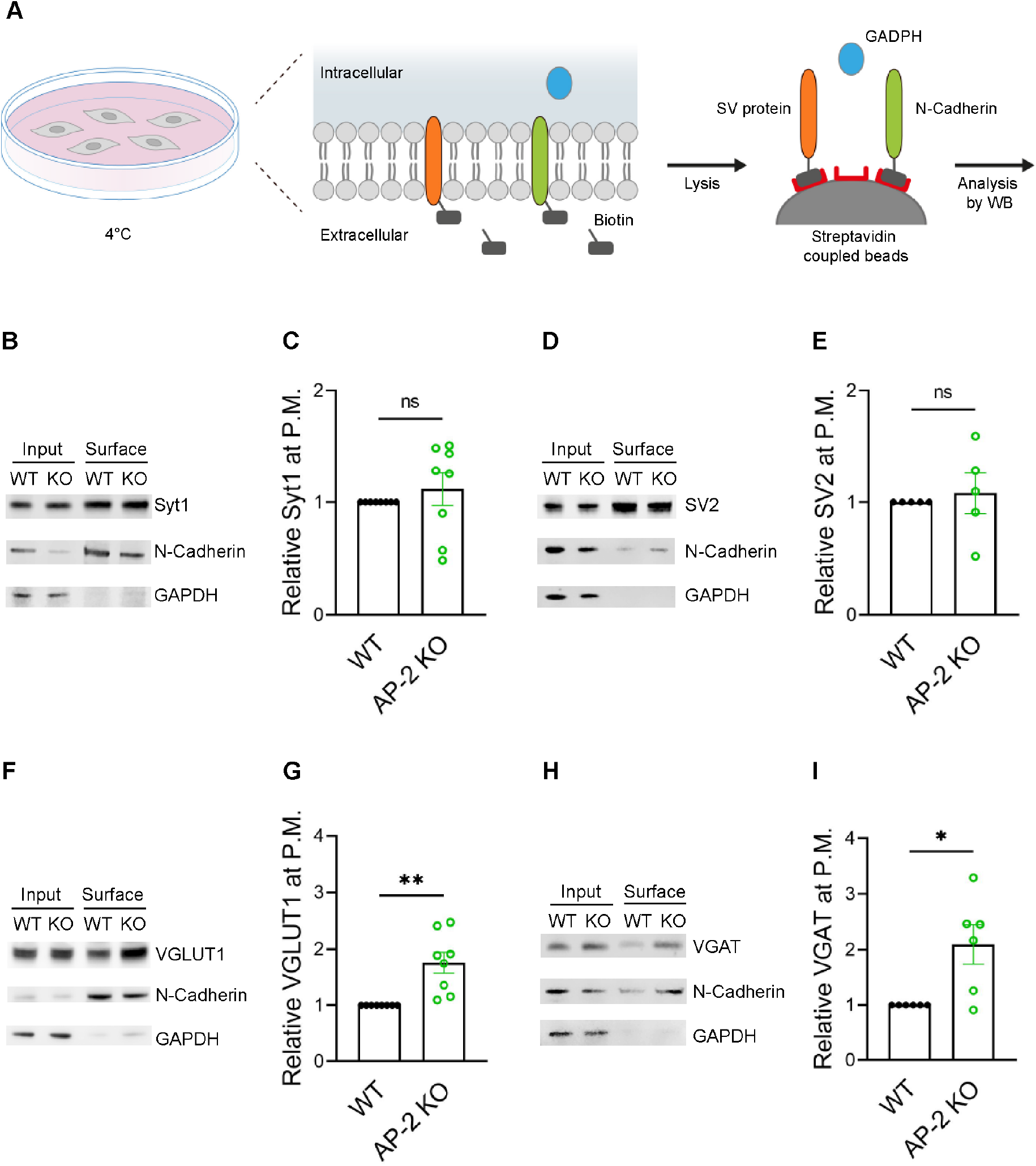
AP-2 depletion results in surface stranding of endogenous vesicular neurotransmitter transporters but not of Synaptotagmin 1 and SV2A. (A) Schematic diagram of the workflow for cell surface protein enrichment. (B-I) AP-2 participates in the surface retrieval of the endogenous SV proteins such as VGLUT1 and VGAT but not of SV2 and Syt1. Cell surface proteins from WT and AP-2 KO cerebellar granule cells were biotinylated and affinity-purified using streptavidin beads. Total (input) and biotinylated proteins (surface) were analyzed by Western blot using antibodies against Syt1 (B), SV2 (D), VGLUT1 (F) and VGAT (H). N-Cadherin and GAPDH were used as control of cell surface membrane and cytosol fraction, respectively. The fold surface enrichment of select proteins (C, E, G, I) in the absence of AP-2 was quantified. Values for WT neurons were set to 1. Data represent the mean ± SEM. n_Syt1_ = 8, n_SV2_ = 5, n_VGLUT1_ = 8, n_VGAT_ = 6 independent experiments. *p*_Syt1_ = 0.4456, *p*_SV2_ = 0.6736, \*\**p*_VGLUT1_ = 0.0049, \**p*_VGAT_ = 0. 0279; two-sided one-sample *t*-test. Raw data can be found in Figure 6-source data 1, and Figure 6-source data 2.

To challenge these results by an independent approach, we took advantage of available antibodies that recognize the luminal domains of VGAT, Syt1 and VGLUT1 (Figure 7). Application of these antibodies under non-permeabilizing conditions to selectively recognize the surface-stranded SV protein pool revealed elevated plasma membrane levels of VGAT (Figure 7F-H) and VGLUT1 (Figure 7-figure supplement 6A-C) in AP-2μ KO hippocampal neurons, while the presynaptic surface pool of Syt1 remained unaltered (Figure 7A-D). Importantly, silencing of neuronal activity in the presence of the sodium channel blocker tetrodotoxin (TTX) rescued surface-stranding of VGAT (Figure 7F-H) and VGLUT1 (Figure 7-figure supplement 6A-C), suggesting that the observed plasma membrane accumulation of a subset of SV proteins in AP-2μ KO neurons is a consequence of defective stimulation-induced SV protein retrieval following exocytic SV fusion.

**Figure 7.**
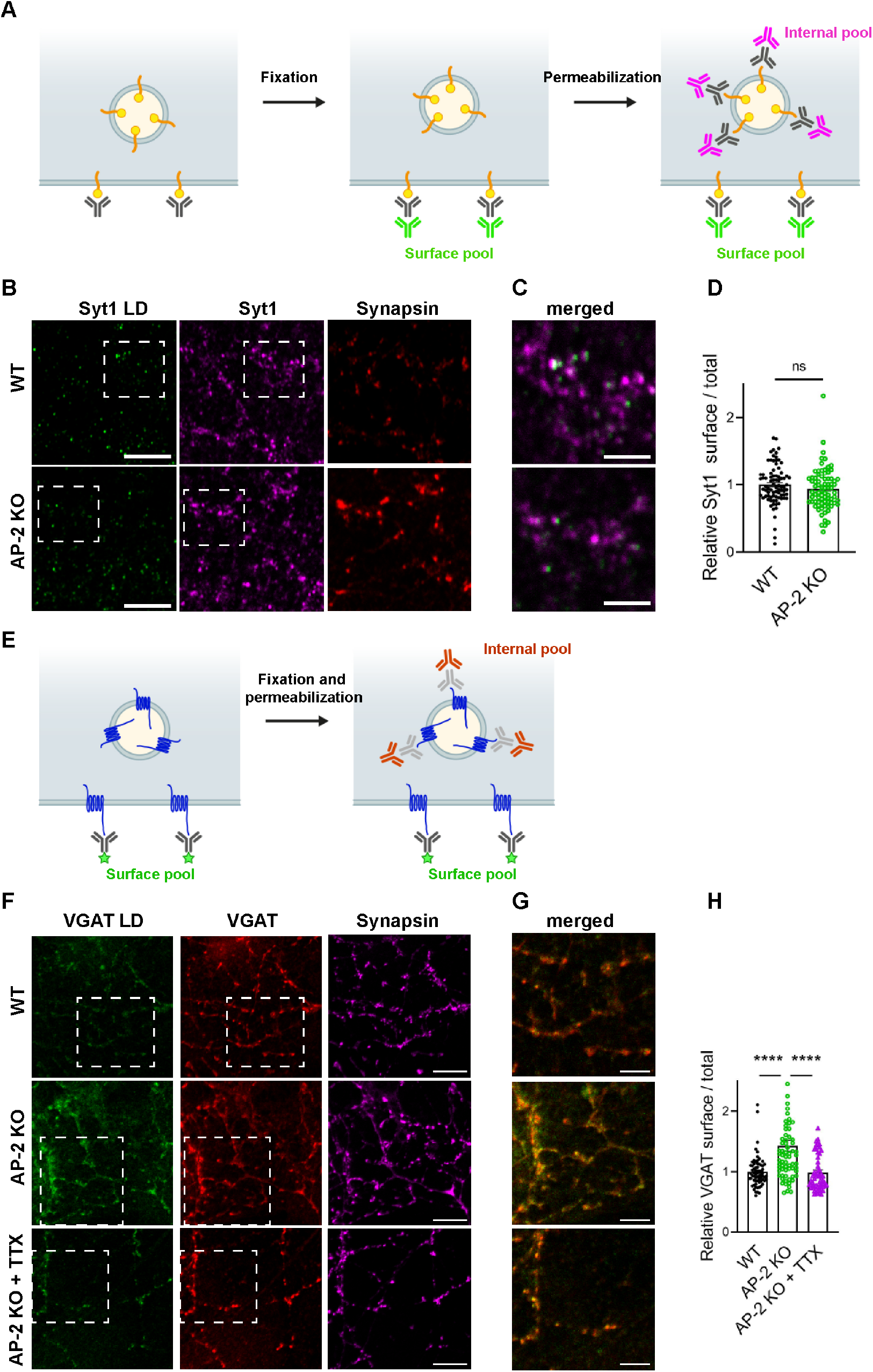
Surface stranding of AP-2-dependent SV cargos in absence of AP-2 is activity-dependent. (A-D) Surface levels of Syt1 are unaffected by loss of AP-2. (A) Schematic drawing of the assay to monitor surface and total levels of Syt1. To label surface epitopes of Syt1, living hippocampal neurons are first incubated with anti-Syt1 antibodies against the luminal domain (black) at 4°C to limit its endocytosis prior to fixation. With no permeabilization conditions, neurons are incubated with 488-conjugated secondary antibodies (green) allowing to reveal the surface pool of Syt1. After washing off unbound antibodies, coverslips are subsequently immunostained using Syt1 antibodies against the cytosolic side (grey) after applying permeabilization conditions. Incubation with 647-conjugated secondary antibodies (magenta) will reveal the total amount of Syt1. Coverslips will be imaged to determine the amount of surface and total Syt1 labeling present in synapses by additional immunostaining of the presynaptic marker synapsin (not depicted). (B) Representative confocal images of cultured hippocampal neurons from WT or AP-2 KO mice co-immunostained for total Syt1 (magenta), surface Syt1 (Syt1 LD, green) and synapsin (red). Scale bars, 5 µm. (C) A zoom of the marked area in (B). Scale bars, 2 µm. (D) Quantification of surface/total Syt1 levels. Values were normalized for WT. Data represent mean ± SEM of n_WT_ = 83 images, n_AP2 KO_ = 79 images. *p* = 0.1519, two-sided unpaired t-test. (E-H) Elevated surface levels of VGAT in absence of AP-2 is rescued by blocking neuronal network activity. (E) Schematic drawing of the assay to monitor surface and total levels of VGAT. To label the surface pool of VGAT, living hippocampal neurons are first incubated with fluorophore-conjugated (green stars) antibodies (black) against the lumenal domain of VGAT at 4°C prior to fixation. After permeabilization, coverslips are immunostained using VGAT antibodies against the cytosolic side (grey) and 568-conjugated secondary antibodies (orange) revealing the total VGAT. Coverslips will be imaged for analyzing the surface and total VGAT labeling present in synapses by additional immunostaining of the presynaptic marker synapsin (not depicted). (F) Representative confocal images of WT or AP-2 KO hippocampal neurons treated or not with tetrodotoxin (TTX) since DIV7 to block spontaneous action potentials and co-immunostained for total VGAT (red), surface VGAT (VGAT LD, green) and synapsin (magenta). Scale bar, 10 µm. (G) A zoom of the marked area in (F). Scale bars, 5 µm. (H) Quantification shows that elevated ratio of surface/total VGAT in AP-2 KO neurons is rescued when neurons were treated with TTX. Values were normalized to WT. Data represent mean ± SEM of n_WT_ = 67 images, n_AP2 KO_ = 69 images and n_AP2 KO+TTX_ = 58 images. \*\*\*\**p*_WT vs AP2 KO_ < 0.0001, *p*_WT vs AP2 KO+TTX_ = 0.9904, \*\*\*\**p*_AP2 KO vs AP2 KO+TTX_ < 0.0001. One-way ANOVA with Tukey’s post-test. Raw data can be found in Figure 7-source data 1.

Collectively, these findings provide strong support for the hypothesis that the clathrin adaptor AP-2 is required for the endocytic retrieval of select SV cargos including VGLUT1 and VGAT under physiological conditions, thereby identifying a clathrin-independent function of AP-2 in the sorting of SV proteins at the presynaptic plasma membrane at central mammalian synapses.

### AP-2 binding deficient mutations in vesicular transporters phenocopy loss of AP-2

In a final set of experiments, we set out to determine the molecular basis for the clathrin-independent function of AP-2 in the sorting of SV proteins by focusing on VGLUT1 and VGAT. Previous studies had identified acidic cluster dileucine motifs (43, 44) in the cytoplasmic tails of vesicular neurotransmitter transporters as putative interaction sites for AP-2 and possibly other clathrin adaptors (Figure 8A). As mutational inactivation of these motifs was further reported to delay the kinetics of VGLUT1 and VGAT analyzed at non-physiological temperature (26–28), vesicular neurotransmitter transporters were proposed to be internalized via CME mediated by clathrin and AP-2 (33). Given our data reported above, we hypothesized that these prior results might reflect the direct recognition of VGLUT1 and VGAT by AP-2 at the neuronal surface to enable their internalization via CIE at physiological temperature (see Figure 1C).

**Figure 8.**
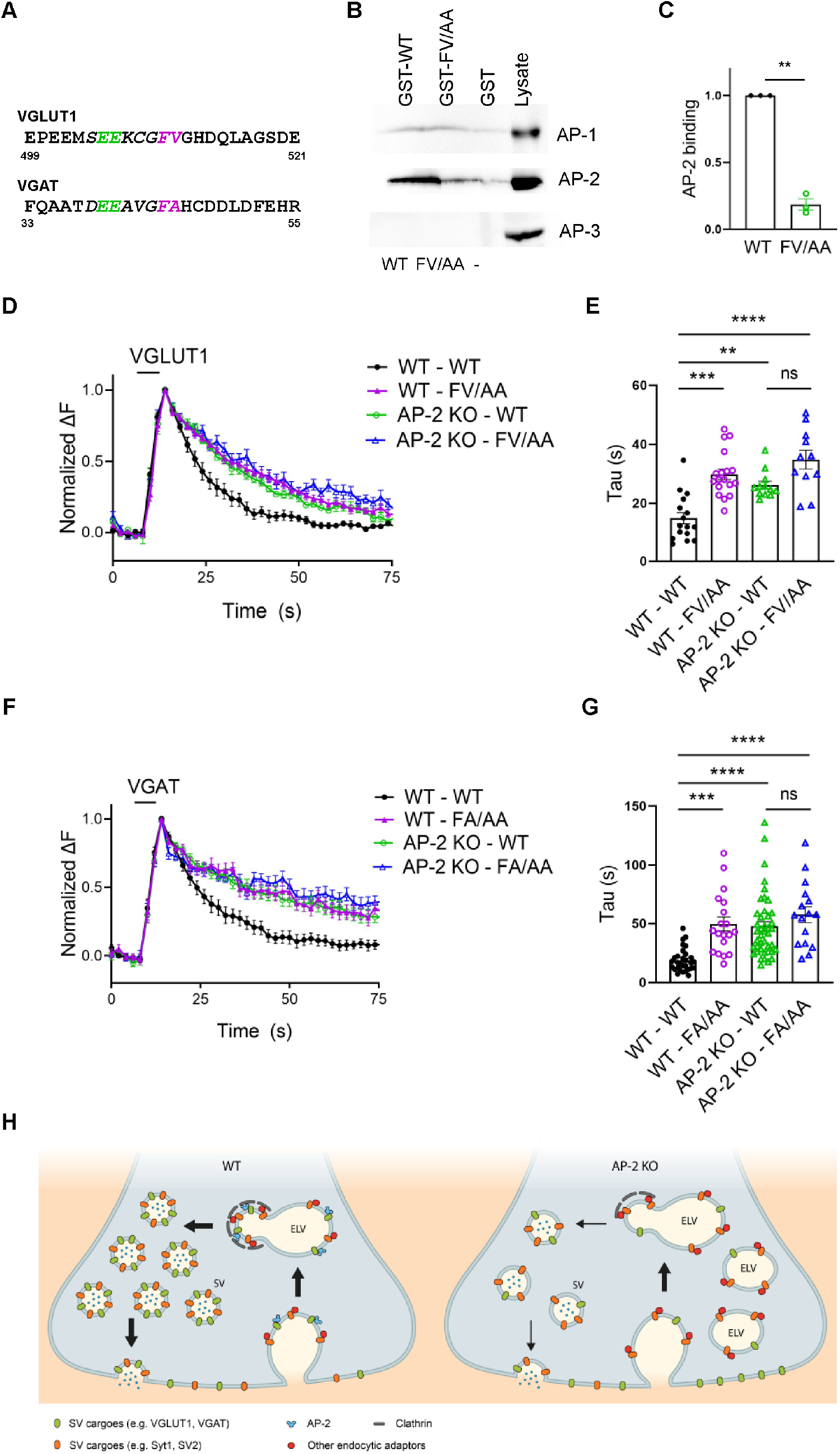
AP-2 binding deficient mutants of vesicular neurotransmitter transporters phenocopy loss of AP-2. (A-C) Association of the cytoplasmic domain of VGLUT1 with the clathrin adaptor complex AP-2 is abolished upon mutational inactivation of the putative AP-2 binding dileucine motif, i.e. F510A/V511A (FV/AA). (A) Acidic cluster di-leucine-like motifs identified in the C-terminal cytoplasmic tail of mouse VGLUT1 and in the N-terminal cytoplasmic tail of mouse VGAT. Numbers indicate amino acid numbers of the respective proteins. Green and magenta indicate two acidic amino acids and two hydrophobic amino acids conserved within the motifs. (B) Immunoblot analysis of material affinity-purified via GST-VGLUT1 C-terminus-WT, GST-VGLUT1 C-terminus-FV/AA or GST alone and brain lysate using specific antibodies against AP-2, AP-1 and AP-3 shows that di-leucine-like motif found in the C-terminus of VGLUT1 binds preferentially to AP-2. (C) Quantified data exhibit VGLUT1 C-terminus-FV/AA variant to significantly disrupt interaction with AP-2. Data represent the mean ± SEM from n = 3 independent experiments. \*\**p* = 0.0027, two-sided one-sample *t*-test. (D-G) Mutant variants of VGLUT1 or VGAT defective in AP-2 binding display significantly slower endocytosis kinetics in response to stimulation in a similar manner to be observed in absence of AP-2. (D) Average normalized traces of neurons from WT and AP-2 KO mice transfected with either the WT or the mutant variant (FV/AA) of VGLUT1-SEP in response of a stimulus train of 200 AP applied at 40 Hz at PT. (E) Quantification of the endocytic decay constant (Tau) of VGLUT1-SEP-expressing neurons (τ_WT-WT_ = 14.82 ± 1.977 s, τ_WT-FV/AA_ = 29.82 ± 1.815 s, τ_AP2 KO-WT_ = 26.15 ± 1.342 s, τ_AP-2 KO-FV/AA_ = 34.80 ± 3.178 s). Data represent the mean ± SEM of n_WT-WT_ = 16 images, n_WT-FV/AA_ = 18 images, n_AP2 KO-WT_ = 12 images, n_AP2 KO-FV/AA_ = 11 images. \*\*\*\**p*_WT-WT vs WT-FV/AA_ < 0.0001, \*\**p*_WT-WT vs AP2 KO-WT_ = 0.0023, \*\*\*\**p*_WT-WT vs AP2 KO-FV/AA_ < 0.0001, *p*_AP2 KO-WT vs AP2 KO-FV/AA_= 0.0530, one-way ANOVA with Tukey’s post-test. (F) Average normalized traces of WT and AP-2 KO neurons transfected with either the WT or the mutant variant (FA/AA) of VGAT-SEP in response of 200 AP applied at 40 Hz at PT. (G) Quantification of the endocytic decay constant (Tau) of VGAT-SEP-expressing neurons (τ_WT-WT_ = 18.86 ± 1.789 s, τ_WT-FA/AA_ = 49.56 ± 5.951 s, τ_AP2 KO-WT_ = 47.43 ± 4.194 s, τ_AP2 KO-FA/AA_ = 57.60 ± 6.920 s). Data represent the mean ± SEM of n_WT-WT_ = 30 images, n_WT-FV/AA_ = 19 images, n_AP2 KO-WT_ = 42 images, n_AP2 KO-FA/AA_ = 16 images. \*\*\**p*_WT-WT vs WT-FA/AA_ = 0.0001, \*\*\*\**p*_WT-WT vs AP2 KO-WT_ < 0.0001, \*\*\*\**p*_WT-WT vs AP2 KO-FA/AA_ < 0.0001, *p*_AP2 KO-WT vs AP2 KO-FA/AA_= 0.4561, one-way ANOVA with Tukey’s post-test. (H) Illustrated model proposing a clathrin independent role for dedicated endocytic adaptors such as AP-2 which recognize select exocytosed SV proteins (e.g. VGLUT and VGAT) present on the neuronal surface to facilitate their clathrin-independent endocytic internalization while clathrin operates downstream facilitating the reformation of functional SVs by budding from internal endosome like vacuoles in a process that also depends on AP-2 and other clathrin-associated endocytic proteins. Raw data can be found in Figure 8-source data 1,and Figure 8-source data 2.

To probe this hypothesis, we first analyzed the association of the cytoplasmic C-terminal domain of VGLUT1 with the clathrin adaptor complex AP-2 and its close relatives AP-1 and AP-3. Robust binding of the GST-fused cytoplasmic domain of VGLUT1 to AP-2 was observed, whereas no association with AP-3 was detected (Figure 8B, C). Mutational inactivation of the putative AP-2 binding dileucine motif, i.e. F510A/ V511A (26–28), largely abrogated VGLUT1 complex formation with AP-2. We also detected a weak, possibly non-specific interaction of VGLUT1 with AP-1 that was insensitive to the F510A/ V511A mutation (Figure 8-figure supplement 7A, B). These results show that VGLUT1 is directly recognized and binds to AP-2 via its acidic cluster dileucine motif.

To probe the functional significance of this interaction we monitored the endocytic retrieval of VGLUT1 and VGAT carrying mutations in their respective AP-2 dileucine binding motifs at physiological temperature. Mutant forms of VGLUT1 or VGAT defective in AP-2 binding displayed significantly slower endocytosis kinetics compared to the respective WT proteins (Figure 8D-G, wild-type in black and FV/AA mutant in purple). These endocytic defects were exarcerbated when endocytosis was monitored at room temperature and under conditions that might favor CME (200 APs, 5 Hz; (8)) (Figure 8-figure supplement 7B-E), consistent with earlier data (26). Importantly, the delayed decay of mutant VGLUT1^F510A/V511A^ -SEP signals could not be attributed to defects in reacidification (e.g. caused by internalization into slowly acidifying compartments), because post-stimulus application of acid solution effectively quenched its fluorescence (Figure 8-figure supplement 7F,G). To analyze whether the observed kinetic delay in the endocytosis of dileucine mutant VGLUT1 and VGAT variants was caused by loss of their ability to associate with AP-2, we monitored their retrieval in AP-2μ KO neurons. Strikingly, loss of AP-2 not only phenocopied the effect of mutational inactivation of the dileucine motifs in VGLUT1 or VGAT but combined mutational inactivation of the dileucine motifs in VGLUT1 or VGAT and AP-2μ KO did not result in additive phenotypes (Figure 8D-G).

These data show that AP-2 recognizes surface-stranded VGLUT1 and VGAT via acidic cluster dileucine motifs contained in their cytoplasmic domains to facilitate their endocytic retrieval from the plasma membrane via CIE.

## DISCUSSION

Our findings based on lentiviral depletion of clathrin and conditional KO of AP-2 in hippocampal neurons reveal a crucial clathrin-independent function of the clathrin adaptor AP-2 in the endocytic sorting of a subset of SV proteins at central synapses. Several lines of evidence support this view: First, comprehensive survey of the endocytic retrieval of six major SV proteins by optical imaging conducted in two independent laboratories provides strong support for the emerging notion (9, 11) that SV endocytosis occurs independent of clathrin, corroborating the prevalence of CIE at physiological temperature. Second and most surprisingly, we find that the endocytic retrieval of a subset of these SV proteins including VGLUT1 and VGAT from the presynaptic plasma membrane depends on sorting by the clathrin adaptor AP-2. This conclusion from SEP-based and cypHer5E-based imaging experiments of exogenously expressed or endogenous SV proteins is further corroborated by the observation that a fraction of endogenous VGLUT1 and VGAT molecules remain stranded on the presynaptic plasma membrane of AP-2μ KO neurons, a phenotype that is rescued upon silencing of neuronal activity. Finally, we show that AP-2-mediated efficient sorting of VGLUT1 and VGAT during CIE is achieved by the recognition of acidic cluster dileucine motifs by AP-2, in agreement with earlier biochemical and cell biological experiments (26–28). Our data thus underscore the importance of AP-2-mediated sorting of select SV cargo during CIE, in the absence of which the compositional integrity of SVs becomes perturbed. This mechanism may also be of pathological relevance in humans. For example, defective endocytosis of VGAT and resulting defects in inhibitory neurotransmission may underlie developmental and epileptic encephalopathy caused by a pathogenic loss-of-function variant of AP-2μ in human patients (45).

Our findings are most consistent with and support a mechanism of SV recycling in which dedicated endocytic adaptors such as AP-2 and others (e.g. AP180, Stonin 2) recognize and recruit SV proteins to the site of endocytosis to facilitate their clathrin-independent endocytic internalization via CIE. Clathrin only assembles once endocytic vesicles have pinched off from the plasma membrane, i.e. downstream of CIE, to reform functional SVs by budding from internal ELVs in a process that depends on AP-2 and other clathrin-associated endocytic proteins (Figure 8H). Such an integrated model not only explains previous observations pertaining to the speed of SV endocytosis (7, 10) and the apparent lack of effect of clathrin loss on SV membrane internalization in various models (8, 9, 22, 23), but is also consistent with the slow kinetics of clathrin assembly (46) and the accumulation of SV proteins on the neuronal surface in the absence of dedicated endocytic adaptors for SV proteins, e.g. Stonin 2, AP180/ CALM (3, 30–33), and AP-2 (this study). We speculate that the mechanism identified here also operates during UFE. Interestingly, recent quantitative proteomic analysis of rodent brain has revealed AP-2 but not clathrin to be highly enriched on SVs (47), suggesting that AP-2 may interact with SV cargos prior to the *bona fide* endocytic process. This perpetual interaction of AP-2 with SV cargos might thus enable rapid sorting and endocytic internalization, i.e. during UFE or other forms of CIE. Of note, our findings are also consistent with recent data regarding a calcium-independent form of SV endocytosis that appears to operate independent of clathrin (48).

An important question raised by our work is how AP-2-independent SV cargos such as Synaptotagmin 1 and SV2 are endocytically retrieved from the neuronal surface. While endocytic adaptors for SV2 have not been reported, studies in mouse hippocampal neurons (49, 50) and in invertebrate models (51–53) have identified cargo-selective roles of Stonin 2 and the related SGIP1 protein in the endocytic retrieval of Synaptotagmin 1 from the presynaptic cell surface. Enigmatically however, it was shown that while loss of Stonin 2 causes the partial accumulation of Synaptotagmin 1 at plasma membrane sites near the active zone, the kinetics of SV endocytosis appeared to be even accelerated (31, 49), suggesting a possible function of surface-stranded Synaptotagmin 1 in regulating the speed of SV endocytosis. One possibility therefore is that Synaptotagmin 1, unlike other SV proteins, due to its comparably large presynaptic surface fraction (54) does not require active endocytic sorting, at least during single rounds of calcium-evoked exocytosis and CIE-mediated retrieval. In the course of CIE, significant amounts of membrane are internalized within short time intervals to form ELVs (10). Hence, it is conceivable that Synaptotagmin 1 reaches such sites of endocytic membrane invagination either by lateral diffusion or via confinement near sites of exocytic release (40, 49, 55). The latter may be facilitated by membrane lipids (e.g. cholesterol, phosphatidylinositol 4,5-bisphosphate) and/ or lateral sequestration by endocytic sorting adaptors such as Stonin 2 (49). SV2 may follow Synaptotagmin 1 via a piggy-back mechanism, consistent with the finding that both proteins can form a stable complex *in vivo* and that their endocytic retrieval appears to be coupled (31, 56–58). A confinement-based endocytic mechanism for Synaptotagmin 1/ SV2 sorting is further consistent with the observation that Synaptotagmin 1 and SV2A display comparably little inter-vesicle variation with respect to their copy numbers compared to other SV proteins (59). Future studies are needed to address this intriguing possibility in more detail.

From a more general perspective our findings dissent from the widely held view that AP-2 obligatorily associates with clathrin to execute its cell physiological functions, at least in the central nervous system (CNS) neurons. While the most well-known function of AP-2 is its involvement in CME in mammalian cells and tissues, studies in higher fungi have uncovered a clathrin-independent role of fungal AP-2 in the polar localization of the lipid flippases DnfA and DnfB (60). Interestingly, in this system AP-2 is seen to colocalize with endocytic markers and the actin-associated protein AbpA, but not with clathrin (60). Because AP-2 also colocalizes with a fungal homolog of Synaptobrevin (60), and clathrin-independent SV endocytosis at hippocampal synapses depends on actin polymerization (9), our newly observed function of AP-2 might reflect an unexpectedly widely-conserved endocytic mechanism. Conversely, studies in AP-2 KO cells have revealed AP-2-independent forms of CME in mammals that impact on receptor sorting and signaling (61).

Taken together our findings together with other studies suggest an unexpected plasticity of endocytic mechanisms in eukaryotes including the mammalian CNS.

## MATERIALS AND METHODS

### Animals

Primary neurons for the experiments presented in Figures 2, 4 and the Figure Supplements 1 and 7 were obtained from ICR mice. Pregnant ICR mice were purchased from SLC, Japan. All animal experiments were approved by the Institutional Animal Care and Use Committee of Doshisha University.

Primary neurons for the experiments presented in Figures 3-8 and the Figures Supplements 2-6 were obtained from either wild-type C57BL/6 or conditional AP-2 KO (AP-2^lox/lox^ × inducible CAG-Cre) mice previously described (8). All animal experiments were reviewed and approved by the ethics committee of the “Landesamt für Gesundheit und Soziales” (LAGeSo) Berlin) and were conducted accordingly to the committee’s guidelines.

All mice were given food and water ad libitum. Animals were kept in a local animal facility with a 12-h light and 12-h dark cycle. Ambient temperature was maintained around 21°C with a relative humidity of 50%. The health reports can be provided upon request. Mice from both genders were used for experiments. Littermates were randomly assigned to experimental groups. Multiple independent experiments were carried out using several biological replicates specified in the figure legends.

### Preparation of Neuronal Cell Cultures

Primary hippocampal cultures for the experiments performed in Figures 2, 4 and the Figure Supplements 1 and 7 were prepared from embryonic day 16 ICR mice as described previously (38, 62), with slight modifications. Briefly, hippocampi were dissected, and incubated with papain (90 units/mL, Worthington) for 20 min at 37°C. After digestion, hippocampal cells were plated onto poly-D-lysine-coated coverslips framed in a Nunc™ 4-well dish (Thermo Fisher) at a cell density of 20,000–30,000 cells/cm^2^ and grown in Neurobasal™ medium (Thermo Fisher) supplemented with 1.25% FBS, 2% B27 and 0.5 mM glutamine at 37°C, 5% CO_2_. On 2-3 days *in vitro* (DIV), 40 μM FUDR (Sigma) and 100 μM uridine (Sigma) were added to the culture medium to limit glial proliferation. One-fifth of the culture medium was routinely replaced with fresh Neurobasal™ medium supplemented with 2% B27 and 0.5 mM glutamine every 2-4 days.

To prepare primary hippocampal and cerebellar neurons for the experiments presented in Figures 3-8 and the Figure Supplements 2-6, hippocampus or cerebellum were surgically removed from postnatal mice at P1-3 or P6, respectively. This was followed by trypsin digestion and dissociation into single neurons. Primary neurons were plated onto poly-L-lysine-coated coverslips for 6-well plates and cultured in MEM medium (Thermo Fisher) containing 2% B27 and 5% FCS. The medium for cerebellar cultures additionally contained 25 mM KCl. To avoid astrocyte growth, hippocampal cultures were treated with 2 μM AraC. To deplete AP-2μ subunit, cultured neurons from floxed conditional AP-2 KO mice expressing a tamoxifen-inducible Cre recombinase were treated with 0.25 μM (Z)-4-hydroxytamoxifen (Sigma) at DIV3. Neurons derived from floxed littermates that were Cre negative were used as controls and treated with equal amounts of (Z)-4-hydroxytamoxifen.

### Plasmids

SEP-tagged Syt1 (NM_001252341.1), VGLUT1 (NM_182993.2) and SV2 (NM_057210.2) were designed as previously described (26, 63, 64), and generated by In-Fusion recombination (Takara Bio). VGAT-SEP was constructed by fusing SEP to the luminal C-terminus of VGAT (NM_031782.2), preceded by GAATCC via In-Fusion recombination. Syp-SEP and Syb2-SEP were kind gifts from L. Lagnado (Sussex, UK) and S. Kawaguchi (Kyoto, Japan), respectively (17, 65). All SEP-tagged constructs were cloned in pcDNA3.1 expression vector or pLenti6PW lentiviral expression vector carrying a TRE promoter (66). pLenti6PW lentiviral expression vector containing a human synapsin 1 promoter that drives a neuron-specific expression of advanced tetracycline transactivator (tTAad) was a generous gift from Y. Fukazawa (Fukui, Japan), and used to induce a protein expression under the control of TRE promoter (66). VGLUT1-F_510_V_511_/AA and VGAT-F_44_A/AA were made by PCR mutagenesis. Cytoplasmic C-terminal region of VGLUT1 (a.a. 496-560) was determined using Expasy ProtScale (https://web.expasy.org/protscale/) and subcloned into pGEX6P1 vector via *BamH*I and *Sal*I sites. For experiments presented in Figures 2, 4 and Supplementary Figures 1 and 5, U6-promoter-based lentiviral shRNA vectors targeting mouse CHC (5’-GTTGGTGACCGTTGTTATG-3’) (11) or luciferase (5’-CCTAAGGTTAAGTCGCCCTCG-3’) as a non-silencing control (67) were obtained from VectorBuilder biotechnology Co. Ltd (Kanagawa, Japan). To identify transduced cells, the shRNA vectors contained a mCherry sequence downstream of a hPGK promoter sequence.

For the data presented in Figures 3 and 4, plasmids encoding for Syb2 and Syt1 (68) with a TEV protease cleavable SEP-tag were a kind gift from J. Klingauf (University of Münster, Münster, Germany). Syp-SEP was a kind gift from L. Lagnado (Sussex, UK). VGLUT1-SEP-tag was a kind gift from R. Edwards and S. Voglmaier (UCSF, CA, USA). SV2A-SEP (69) was a kind gift from E.R. Chapman (UW-Madison, WI, USA). VGAT-SEP was a gift from S. Voglmaier (Addgene plasmid #78578; http://n2t.net/addgene:78578 ; RRID:Addgene_78578). For the clathrin knockdown experiments performed in Figure 5 and the Figure Supplement 4, expression vectors f(U6)sNLS-RFPw msClathrin scrambled and f(U6)sNLS-RFPw msClathrin shRNA were a kind gift from C. Rosenmund (Berlin, Germany) (8, 11). For rescue experiments shown in Figure 3, we used a construct previously described (70) containing murine untagged AP-2μ followed by an IRES site by mRFP in an adenoviral AAV-HBA-EWB vector backbone.

### Antibodies

#### Immunoblotting

Secondary antibodies were all species-specific. Horseradish peroxidase (HRP)-conjugated or LI-COR 800CW and 680RD infrared suitable antibodies were applied at 1:10.000 in blocking solution. Quantification was done based on chemiluminescence or fluorescence using an Odyssey FC detection system. Each panel of a figure has individual antibodies shown at the same exposure settings throughout the experiment.

#### Immunofluorescence

Secondary antibodies were all species-specific. Secondary antibodies fluorescently labelled with Alexa dyes 488, 568 or 647 (Thermo Fisher Scientific) were applied at 1:1000 or 1:500 in blocking solution.

Antibodies used in this study are listed in Supplementary Table 1.

### Drug application

Pitstop 2 (Leibniz-Forschungsinstitut für Molekulare Pharmakologie, Berlin, Germany) was used to inhibit clathrin-dependent endocytosis (39). Growth media of primary hippocampal neurons at DIV13-15 were replaced by osmolarity-adjusted, serum-free NBA medium (Gibco) for 1 h prior the incubation with 30 μM of pitstop 2 during 1 h. After treatment with Pitstop 2, experiments of transferrin uptake or cypHer-labeled antibody-based live imaging were performed. Tetrodotoxin (TTX) (Sigma) was used to inhibit voltage-gated sodium channels and silence synaptic activity in cultured hippocampal neurons. Where indicated, neurons were incubated with 1 μM TTX at DIV7, which was renewed on DIV11.

### Transfection of primary neurons

Calcium phosphate transfection was carried out as previously described (71), with slight modifications, using CalPhos™ Mammalian Transfection Kit (Takara Bio or Promega) on DIV7. Shortly, 6 μg plasmid DNA and 248 mM CaCl_2_ dissolved in water were mixed with equal volume of 2x HEPES buffered saline (total of 100 μl for one 35 mm dish), and incubated for 15-20 min allowing for precipitate formation, while neurons were starved in osmolarity-adjusted, serum-free MEM (Sigma) or NBA medium (Gibco) for the same time at 37°C, 5% CO_2_. Precipitates were added to neurons and incubated at 37°C, 5% CO_2_ for 30-40 min. Finally, neurons were washed by incubation in fresh MEM or osmolarity-adjusted HBSS (Gibco) at 37°C, 10% CO_2_ for 15 min and transferred back into their conditioned medium. For rescue experiments AP-2μ-IRES-RFP construct was introduced at DIV7 and the neurons were analyzed at DIV14.

### Lentivirus transduction of primary neurons

For experiments presented in Figures 2, 4 and the Figure Supplements 1 and 7, lentivirus was produced as described previously (38, 62). The cells were transduced with lentivirus expressing VGLUT1-SEP and its mutant on DIV2, Syt1-SEP, Syp-SEP, Syb2-SEP, VGAT-SEP and its mutant on DIV5-7, and shRNA for CHC on DIV7. To activate protein expression under the control of TRE promoter, lentivirus expressing tTAad was co-transduced on the same DIV. Transduction of clathrin shRNA on earlier DIV caused severe loss of neurons at the time of recordings, and thus, cells were only transduced with clathrin shRNA on DIV7 and all the clathrin knockdown experiments were performed on DIV14. For the clathrin knockdown experiments, lentiviral particles were prepared as follows: HEK293T cells were co-transfected with the lentivirus shuttle vector (10 μg) and two helper plasmids, pCMVdR8.9 and pVSV.G (5 μ g each) using the calcium phosphate method. After 48 and 72 hours, virus-containing supernatant was collected, filtered, aliquoted, snap-frozen in liquid nitrogen and stored at −80°C. Viruses were titrated with mice WT hippocampal mass-cultured neurons using NLS-RFP signals. For the clathrin knockdown experiments performed in Figure 5 and the Figure Supplement 4 mouse hippocampal neurons were transduced at DIV2, resulting in clathrin heavy chain depletion at DIV14 from the start of the treatment.

### Live Imaging

For experiments presented in Figures 2, 4 and the Figure Supplements 1 and 7, fluorescence imaging was performed on IX71 inverted microscope (Olympus) equipped with a 60x (1.35 NA) oil immersion objective and 75-W xenon arc lamp (Ushio). Cells on coverslips were mounted on a custom-made imaging chamber equipped on a movable stage with constant perfusion of Tyrode’s solution (140 mM NaCl, 2.4 mM KCl, 10 mM HEPES, 10 mM glucose, 2 mM CaCl_2_, 1 mM MgCl_2_, 0.02 mM CNQX, 0.025 mM D-APV, adjusted to pH 7.4). Temperature was clamped at physiological temperature (PT) (35 ± 2°C) or room temperature (RT) (25 ± 2°C) using TC-324C temperature controller (Warner Instruments) with feedback control, SH-27B in-line solution heater (Warner Instruments) and a custom-equipped air-heater throughout the experiment. Electrical field stimulation was delivered via bipolar platinum electrodes with 1-ms constant voltage pulses (50 V) controlled by pCLAMP™ Software (Molecular Devices). Fluorescence images (512 × 512 pixels) were acquired with ORCA-Flash 4.0 sCMOS camera (Hamamatsu Photonics) in time-lapse mode either at 1 fps (for imaging in response to 200-APs stimulation) or 2 fps (for imaging in response to 50-APs stimulation) under the control of MetaMorph software (Molecular Devices). SEP fluorescence was imaged with 470/22 nm excitation and 514/30 nm emission filters, and mCherry fluorescence with 556/20 nm excitation and 600/25 nm emission filters.

For SEP-based assays presented in Figures 3, 4, 8 and the Figure Supplement 3 cultured neurons at DIV13-15 were placed into an RC-47FSLP stimulation chamber (Warner Instruments) for electrical field stimulation and imaged at 37°C in osmolarity-adjusted basic imaging buffer (170 mM NaCl, 3.5 mM KCl, 20 mM N-Tris[hydroxyl-methyl]-methyl-2-aminoethane-sulphonic acid (TES), 0.4 mM KH_2_PO_4_, 5 mM glucose, 5 mM NaHCO_3_,1.2 mM MgCl_2_, 1.2 mMNa_2_SO_4_,1.3 mM CaCl_2_, 50 mM AP5 and 10 mM CNQX, pH 7.4) by epifluorescence microscopy [Nikon Eclipse Ti by MicroManager 4.11, eGFP filter set F36-526 and a sCMOS camera (Neo, Andor) equipped with a 40x oil-immersion objective]. For evaluation of general exo-/endocytosis, neurons were stimulated with 200 or 50 APs (40 Hz or 20 Hz respectively, 100 mA) and imaged at 1 fps or 0.5 fps with 100 ms excitation at 488 nm. For quantification of active synapses, signals > 4x S.D. of the noise were considered as the threshold to identify ROIs that show stimulus-dependent changes in SEP-fluorescence signals following stimulation.

To monitor recycling of an endogenous SV protein, labelling of hippocampal neurons with a cypHer-conjugated antibody directed against the luminal domain of VGAT (#131103CpH; Synaptic Systems) was performed by incubating primary neurons with anti-VGAT cypHer at 1:120 (from a 1 mg/ml stock) in their own conditioned culture medium for 1 h. Neurons were then washed with imaging buffer and placed in the stimulation chamber for electrical field stimulation and live imaging was performed as described above for SEP-based assays. Images were acquired at 1fps with 100 ms excitation at 647 nm. For Supplementary figure 3I,J, time-lapse mode was done at 1 frame every 3 seconds.

### Image and Data Analysis

Quantitative analysis of responding boutons was performed in Fiji (72) using Time Series Analyzer plugin (https://imagej.nih.gov/ij/plugins/time-series.html) or by using custom-written macros (available at https://github.com/DennisVoll/pHluorin_ROI_selector/). Circular regions of interest (ROIs, 4-µm^2^ area) were manually positioned at the center of fluorescent puncta that appear stable throughout all trials and responded to an electrical stimulus, and the fluorescence was measured over time. Another five ROIs of the same size were positioned at the regions where no cell structures were visible, and their average fluorescence was subtracted as background signals. After further subtracting base signals, the fluorescence of each time point was normalized with the peak value. Time constant of endocytosis (Tau) was determined by fitting mono-exponential decay curve [0+A*exp(-x/tau)] using ‘scipy.optimize.curve_fit’ function in Python (73) or Prism 8 (Graphpad) softwares. Data of < 30 boutons from a single experiment were averaged and counted as n = 1 for Tau calculation. All data were collected from 2 to 5 independent preparations.

#### Photobleaching correction

Decrease in the fluorescence intensity signals due to photobleaching was corrected as previously described (40). The decay constant τ was obtained from observations of intensity-time courses of non-active boutons by experimentally fitting a monoexponential decay curve as follows: I(t)= A*(exp(-t/τ), with I(t), as fluorescence intensity at time t; A, as initial intensity I(0); and τ, as time constant. The fluorescence intensities following the time course of photobleaching were calculated experimentally for every time point of the recording and summed up to the mean fluorescence intensity at time t.

### Immunocytochemical analysis of cultured neurons

For the experiments presented in the Figure Supplement 1, cultured hippocampal neurons transduced with non-silencing shRNA or shRNA targeting mouse CHC were fixed on DIV14 with 4% (w/v) paraformaldehyde and 4% sucrose in PBS for 15 min at RT. After washing in PBS, fixed cells were permeabilized with 0.2% Triton X-100 in PBS for 10-15 min and blocked in PBS containing 10% (v/v) FBS for 30 min at RT. Cells were then incubated with rabbit anti-CHC (1:1,000, abcam, ab21679) and mouse anti-synaptophysin (1:1,000, a kind gift from R. Jahn [Göttingen, Germany]) antibodies for 2h at RT, and subsequently, with anti-rabbit IgG Alexa Fluor 647 (1:1,000, Thermo Fisher, A-21245) and anti-mouse IgG Alexa Fluor 488 (1:1,000, Thermo Fisher, A-11029) antibodies for 45 min at RT. Transduced cells, visible by mCherry expression, were imaged using the same microscope setup with live imaging. Synaptophysin signals were imaged with 482.5/12.5 nm excitation and 530/20 nm emission filters, CHC signals with 628/20 nm excitation and 692/20 nm emission filters, and mCherry fluorescence with 540/10 nm excitation and 575IF-emission filters. Ratiometric quantification of CHC signals over synaptophysin signals was conducted in the automated fashion using Fiji with a custom-written macro as previously described (10), with slight modifications. In short, acquired images were first background-subtracted using ‘Rolling Ball’ function with the radius set at 30 pixels (http://fiji.sc/Rolling_Ball_Background_Subtraction). Then, the synapses are defined by thresholding the synaptophysin signals using built-in ‘Default’ method (https://imagej.net/Auto_Threshold.html#Default). The binary image of synaptophysin was used as the regions of interest. The average intensities of CHC and synaptophysin were measured from those locations and were divided to obtain ratio between those two proteins. For experiments presented in the Figure Supplements 2 and 4, primary hippocampal neurons seeded on coverslips were fixed for 13 min with 4% PFA in PBS solution on ice and washed three times with PBS. Cells were permeabilized and blocked in blocking solution (PBS, 10% goat serum and 0.1% Triton X-100) for 30 min and incubated with primary antibodies diluted in blocking solution for 1 h. After three washes with PBS, coverslips were incubated for 1 h with secondary antibodies diluted in blocking solution, followed by three washes in PBS. Coverslips were mounted in Immu-Mount (Thermo Fisher) with 1.5 mg ml^−1^ 4,6-diamidino-2-phenylindole (DAPI; Sigma) to stain nuclei and were visualized routinely using the Zeiss laser scanning confocal microscope LSM710.

To distinguish between surface and internal SV protein pool, hippocampal neurons (DIV13-15) were gently washed once using osmolarity-adjusted HBS (25 mM HEPES, 140 mM NaCl, 5 mM KCl, 1.8 mM CaCl_2_, 0.8 mM MgCl_2_, 10 mM glucose, pH 7.4) prior to live labeling surface-localized VGAT, Syt1 or VGLUT1 with specific antibodies against their luminal/ extracellular regions (VGAT Oyster 488: SySy, #131103C2 or VGAT Oyster 568, SySy, 131103C3; Synaptotagmin 1: SySy, #105102; VGLUT1: SySy, #135304) for 20 min at 4°C. After washing twice, cells were fixed for 5 min at RT in 4% PFA and washed again three times for 3 min. Without permeabilization, neurons were stained with Alexa Fluor 488 secondary antibody (except for VGAT which was previously incubated with an Oyster-labeled antibodies) for 45 min at room temperature, to visualize the surface pool of the corresponding SV protein. For labelling the internal population of those SV proteins in the same experiment, neurons were permeabilized for 10 min using 0.1% Triton X-100 and subsequently coverslips were incubated for 1 h at RT with primary antibodies recognizing the cytosolic side of Synaptotamin 1 (SySy, 105 011), VGAT (SySy, 131 013) and VGLUT1 (SySy, #135304); and stained with Alexa Fluor 647 or 568 for 45 min at RT revealing the internal fraction of the SV protein pool. After four final washing steps of 3 min, the coverslips were mounted in Immu-Mount with DAPI to stain nuclei. Samples were visualized using the Zeiss laser scanning confocal microscope LSM710 using a 63x oil objective. All acquisition settings were set equally for all groups within each experiment. Surface and internal fluorescent intensities were individually quantified using ImageJ and the ratio between such surface and internal signals was calculated for WT and KO conditions.

For quantifying the levels of presynaptic proteins (VGAT, VGLUT1 and Syt1) in axons, synapsin staining signals were used as a mask to restrict the quantified are to the shape of synapsin-positive boutons by applying thresholding using ImageJ. Values were normalized to WT.

### Transferrin uptake

Primary neurons expressing lentivirally delivered clathrin heavy chain (CHC)-targeting shRNA or a scramble version (DIV14) were starved for 1 hour in osmolarity-adjusted NBA medium (Gibco) at 37°C, 5% CO_2_ and treated with 25 μg/ml transferrin coupled to Alexa Fluor 647 (Tf-647, Life technologies) in NBA medium for 20 min at 37°C, 5% CO_2_. To remove unbound Tf-647, neurons were washed twice with cold PBS, followed by 1 min of acid-wash at pH 5.3 (cold 0.1 M acetic acid supplemented with 0.2 M NaCl) to quench surface bound Tf-647 and finally twice with cold PBS prior to 30 min fixation at room temperature with 4% (w/v) PFA and 4% sucrose in PBS. Coverslips were mounted in Immu-Mount with DAPI to stain nuclei and were visualized routinely using the Zeiss laser scanning confocal microscope LSM710 and fluorescence intensities per cell were quantified using ImageJ.

### Protein Expression and Purification

GST-fusion proteins were expressed in *Escherichia coli* BL21 cells overnight at 25°C after induction at OD_600_ 0.4-0.7 with 1 mM isopropyl β-D-1-thiogalactopyranoside. Cells were harvested, resuspended in sonication buffer (50 mM Tris-Cl pH 8.0, 50 mM NaCl, 1 mM EDTA), lysed by ultrasonication followed by 1% Triton X-100 treatment and spun at 39,191x g for 30 min at 4 °C. Proteins were then purified from the supernatant using Glutathione Sepharose 4B resin (Cytiva), according to the manufacturer’s instructions, and dialyzed against pull-down buffer (20 mM HEPES pH 7.4, 140 mM NaCl, 1 mM EDTA, 1 mM DTT).

### GST Pull-Down Assay

Extracts from mouse brain were solubilized in pull-down buffer containing cOmplete™ Protease Inhibitor Cocktail (Roche) and 0.1% saponin for 45 min at 4°C, sedimented at 20,400x g for 25 min at 4°C, and the supernatant (∼16 mg total protein) was incubated with 600 μg of GST fusion proteins immobilized on Glutathione Sepharose for 160 min at 4°C. After pelleting, the beads were washed and bound protein was detected by immunoblot analysis using mouse monoclonal anti-γ-adaptin 1 (1:500), anti-α-adaptin 2 (1:100) and anti-σ-adaptin 3 (1:250) antibodies as primary antibodies. Fiji was used to quantify the intensity of bands.

### Cell Surface Biotinylation Assay

Primary cerebellar granule neurons at DIV10 were treated as previously described (70). Briefly, neurons were placed on ice, washed twice with ice-cold PBS^2+^ (137 mM NaCl, 2.7 mM KCl, 8.1 mM Na_2_HPO_4_, 0.5 mM CaCl_2_, 1 mM MgCl_2_, pH 7.4) and incubated with 0.5 mg ml^−1^ sulfo-NHS-LC-biotin (EZ-Link, Pierce/Thermo Scientific) in PBS^2+^ while shaking for 20 min at 4 °C. The biotinylation solution was removed, and surplus biotin was quenched by two 5-min washes with 50 mM glycine in PBS^2+^ at 4 °C on a shaker. Cells were then washed briefly with PBS and scraped into lysis buffer (20 mM HEPES pH 7.4, 100 mM KCl, 2 mM MgCl_2_, 2 mM PMSF, 1% Triton X-100, 0.6% protease inhibitor cocktail (Sigma)). Lysates were incubated on a rotating wheel at 4 °C for 30 min, followed by centrifugation at 17,000 × *g* for 10 min at 4 °C. The protein concentration of the supernatant was determined using a Bradford or BCA assay. Biotinylated molecules were isolated by a 1.5-h incubation of protein samples (between 500-1,000 μg) with streptavidin beads on a rotating wheel at 4 °C. After centrifugation at 3,500 × *g*, the supernatant was transferred to a fresh tube. Beads were extensively washed, and bound protein was eluted with Laemmli buffer with fresh 5% β-mercaptoethanol by heating to 65 °C for 15 min. Equal protein amounts of lysates were separated by SDS–PAGE and analysed by immunoblotting. Bound primary antibodies were detected by incubation with IRDye 680/800CW-conjugated secondary antibodies or alternatively, HRP-conjugated secondary antibodies and ECL substrate (Pierce 32106). Immunoblots were imaged by LI-COR-Odyssey FC detection with Image Studio Lite Version 4.0. N-cadherin and GAPDH were used as markers for the membrane and cytosol fraction, respectively. All experiments were performed at least four times.

### Analysis of primary neuronal culture extracts

Primary neurons expressing lentivirally delivered either clathrin heavy chain (CHC)-targeting shRNA or its inactive scramble version were harvested at DIV14, lysed using RIPA buffer (150 mM NaCl, 1% NP-40 al 1%, 0.5% sodium deoxycholate, 0.1% SDS, 50 mM Tris-HCl, protease inhibitor cocktail (Sigma) and phosphatase inhibitor cocktail (Sigma)). Alternatively, primary neurons derived from WT or AP-2 KO neurons (DIV 14) were lysed in HEPES lysis buffer (20 mM HEPES pH 7.4, 100 mM KCl, 2 mM MgCl_2_, 2 mM PMSF, 1% Triton X-100, 0.6% protease inhibitor cocktail (Sigma)). Lysates were incubated on a rotating wheel at 4 °C for 30 min, followed by centrifugation at 17,000 × *g* for 10 min at 4 °C. The protein concentration of the supernatant was determined using a Bradford or BCA assay. Protein samples were denatured and separated in 10% SDS/PAGE followed by Western blotting using standard procedures, followed by detection with secondary antibodies coupled to horseradish peroxidase and ECL substrate or IRDye 680/800CW-conjugated secondary antibodies. Immunoblots were imaged and quantified by LI-COR-Odyssey FC detection with Image Studio Lite Version 4.0.

### Statistics

Values are depicted as the mean ± standard error of the mean (SEM) as indicated in the figure legends. For comparisons between two experimental groups, statistical significance of normally distributed data was analysed by two-sample, two-sided unpaired Student’s *t*-tests. For comparisons between more than two experimental groups, statistical significance of normally distributed data was analysed by one-way analysis of variance (ANOVA) with a post-hoc test such as the Tukey post-hoc test (see figure legends). One-sample, two-sided *t*-tests were used for comparisons with control group values that had been set to 1 for normalization purposes and therefore did not fulfil the requirement of two-sample *t*-tests or one-way ANOVA concerning the homogeneity of variances. GraphPad Prism v.8 software was used for statistical analysis. The level of significance is indicated in the figures by asterisks (\**p* ≤ 0.05, \*\**p* ≤ 0.01, \*\*\**p* ≤ 0.001, \*\*\*\**p* ≤ 0.0001) and provided in the figure legends as exact *p* values as obtained by the indicated statistic test. No statistical method was used to predetermine sample sizes as sample sizes were not chosen based on a prespecified effect size. Instead, multiple independent experiments were carried out using several sample replicates as detailed in the figure legends. Whenever possible, data were evaluated in a blinded manner.

### Data availability

All data generated and analyzed in this study are included in the manuscript and supporting files. Numerical data as well as raw images of western blots are included in a single Source Data File.

### Contact for Reagent and Resource Sharing

Further information and requests for resources and reagents should be directed to and will be fulfilled by the corresponding contacts V.H. and S.T.

## ACKNOWLEDGEMENTS

This work was supported by grants from JSPS KAKENHI (19H03330), the JSPS core-to-core program A. Advanced Research Networks (grant number: JPJSCCA20170008), a grant from the Takeda Science Foundation to ST, and grants from the Deutsche Forschungsgemeinschaft (SFB958/ TP A01; HA2686/13-1/ Reinhart-Koselleck Program) to VH. We are indebted to Sabine Hahn, Delia Löwe and Silke Zillmann for expert technical assistance with genotyping and preparation of neuronal cultures. We thank Dr. Barth Rossum for his contribution to the illustrations.

## AUTHOR CONTRIBUTIONS

T.L.H., and K.T. performed SEP-based imaging experiments. T.L.H. conducted all experiments involving neurons from AP-2 KO mice and WT littermates, cypHer-based imaging experiments, transferrin uptakes and the detection of surface SV proteins either by biotinylation assays or immunostainings. K.T. analyzed the association and of VGAT and VGLUT1 with adaptor complexes. Y.M. assisted the experimentation at the initial stage of the work. P.K. and S.N. assisted biochemical assays to validate binding of adaptor proteins to the dileucine motif of VGLUT1. T.L.H., K.T., V.H., and S.T. designed and conceptualized the study and wrote the paper with input from all authors.

## FIGURE SUPPLEMENTS

**Figure 2–figure supplement 1.**
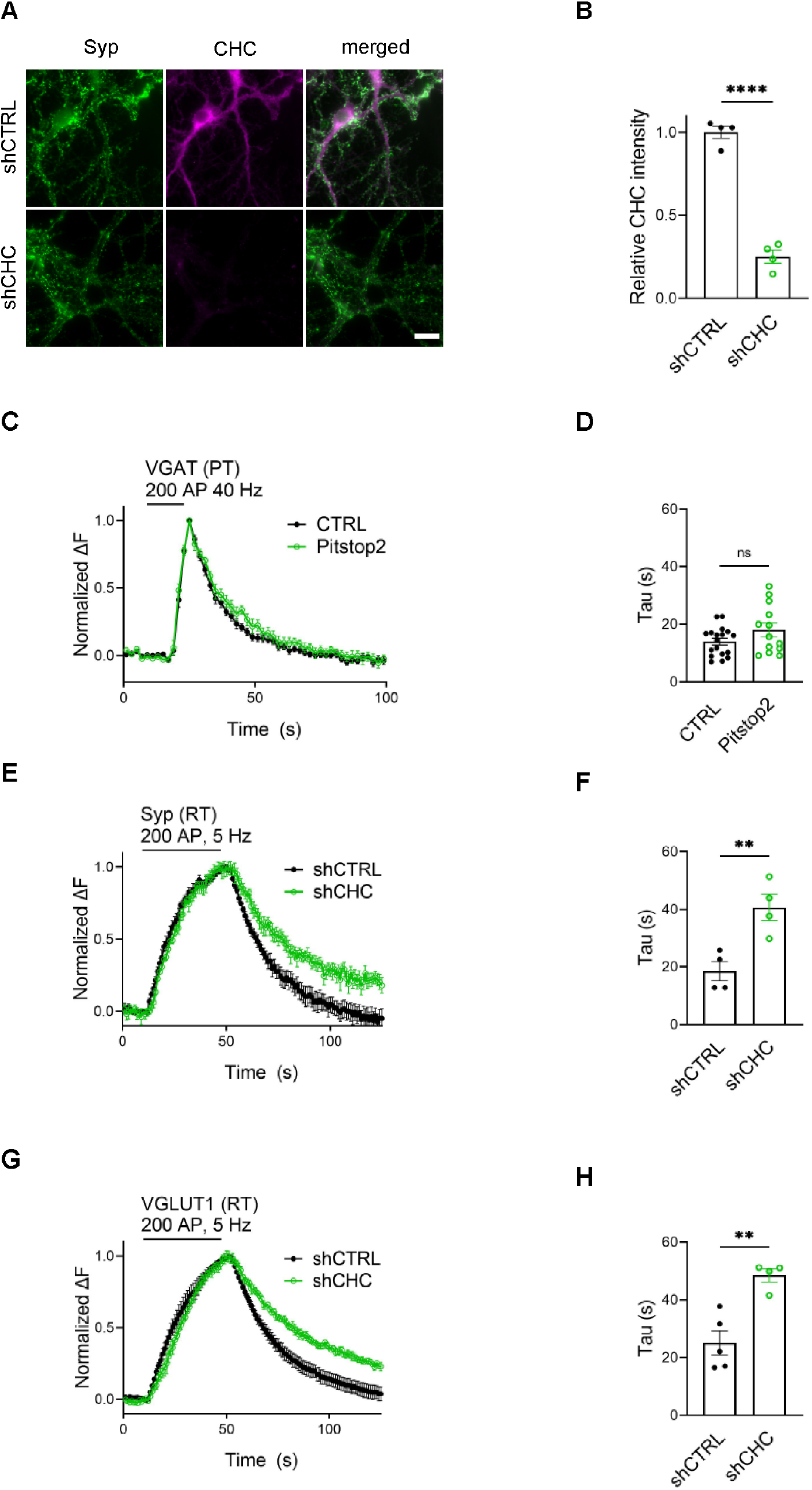
Temperature-sensitive, clathrin-independent SV endocytosis at hippocampal synapses. (A,B) shRNA-targeting clathrin heavy chain (CHC) efficiently depletes CHC in hippocampal neurons (A) Representative images of neurons transduced with lentivirus expressing either non-specific shRNA (shCTRL) or shRNA targeting CHC (shCHC) immunostained for CHC (magenta) and the presynaptic marker synaptophysin (Syp) (green). The scale bar represents 10 μm. (B) Quantification of CHC fluorescence intensity relative to Syp fluorescence. Values for shCTRL were set to 1. Data represent mean ± SEM of n = 4 images for shCTRL and n = 4 images for shCHC. Two-sided unpaired *t*-test, **** *p* < 0.0001. (C,D) Endocytosis of VGAT upon acute inactivation of clathrin by Pitstop 2 proceeds unaffected at PT. (C) Average normalized traces of neurons transfected with VGAT-SEP and treated either with DMSO (CTRL) or Pitstop 2 in response to 200 AP applied at 40 Hz. (D) Endocytic decay constant (Tau) of neurons expressing VGAT-SEP (τ_CTRL_ =14.03 ± 1.159 s, τ_Pitstop2_ = 18.11 ±. 2.315 s). Data shown represent the mean ± SEM with n = 18 images and n=13 images for CTRL and Pitstop 2, respectively. *p* = 0.0976. Two-sided unpaired *t*-test. (E-H) Membrane retrieval induced by low-frequency stimulation at room temperature (RT) is sensitive to clathrin loss. Average normalized traces of neurons transduced with lentivirus expressing shCTRL or shCHC and co-transfected with either Syp-SEP (E) or VGLUT1-SEP (G) upon stimulation of 200 APs at 5 Hz at RT, a condition known to favor CME. Quantification of fluorescence decay (Tau) in neurons co-expressing Syp-SEP (F) and shCTRL (18.5 ± 3.34 s) or shCHC (40.7 ± 4.61 s); and VGLUT1-SEP (H) and shCTRL (25.1 ± 4.18 s) or shCHC (48.5 ± 2.31 s). Data represent the mean ± SEM for Syp-SEP (n_shCTRL_ = 4 images, n_shCHC_ = 4 images; **p = 0.009) and for VGLUT1-SEP (n_shCTRL_ = 5 images, n_shCHC_ = 4 images; \*\**p* = 0.003). Two-sided unpaired *t*-test. Raw data can be found in Figure 2-figure supplement 1-source data 1.

**Figure 3–figure supplement 2.**
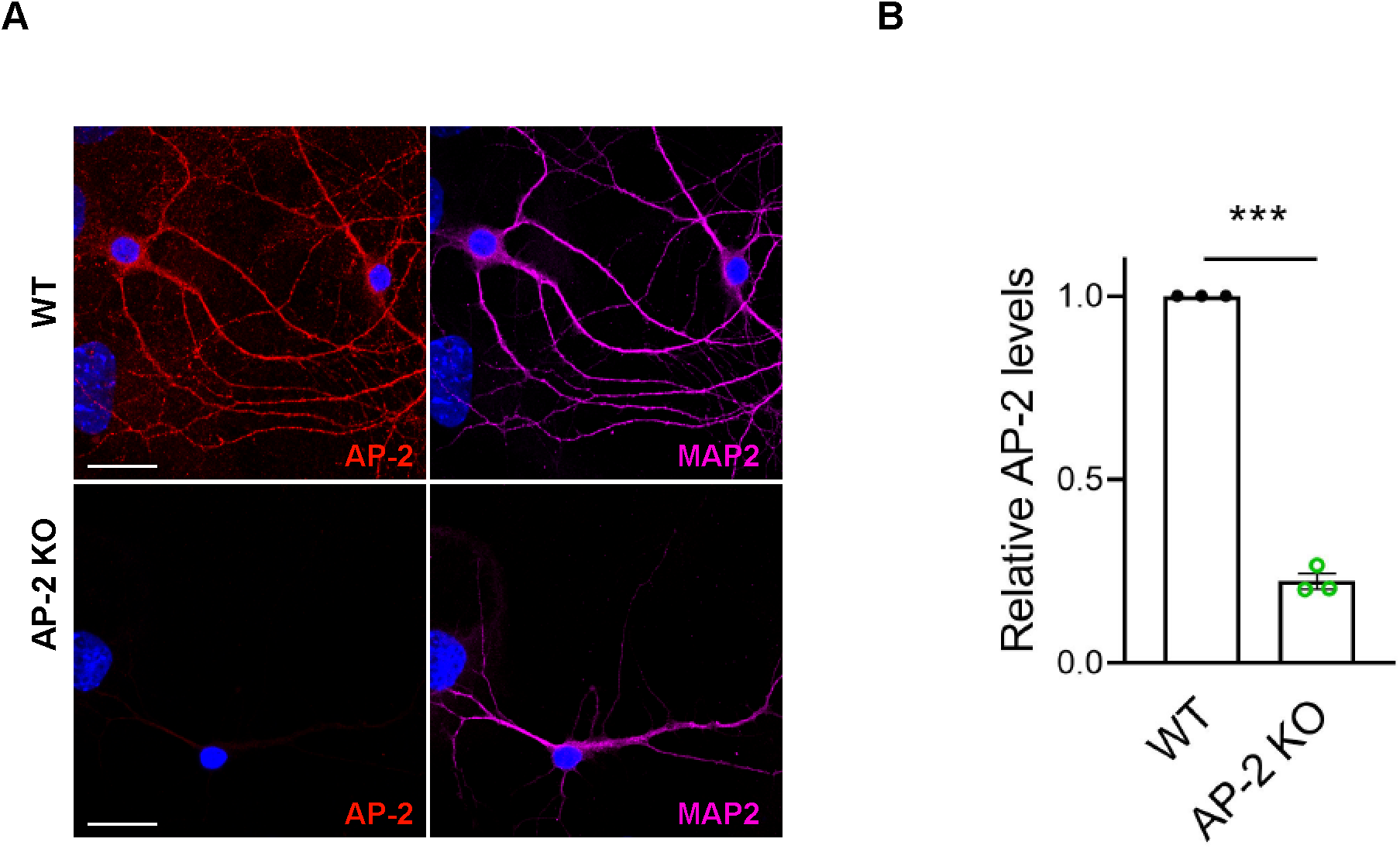
AP-2 depletion in hippocampal neurons. (A) Representative images of WT and AP-2 KO neurons immunostained for AP-2 (red) and the neuronal marker MAP2 (magenta). The scale bar represents 20 μm. (B) Quantification of AP-2 fluorescence intensity. Values for WT neurons were set to 1. Data represent mean ± SEM of n = 3 independent experiments with n = 9 images for WT and n = 8 images for AP-2 KO neurons; \*\*\**p* = 0.0008. *p* = 0.4267, two-sided unpaired *t*-test. Raw data can be found in Figure 3-figure supplement 2-source data 1.

**Figure 4–figure supplement 3.**
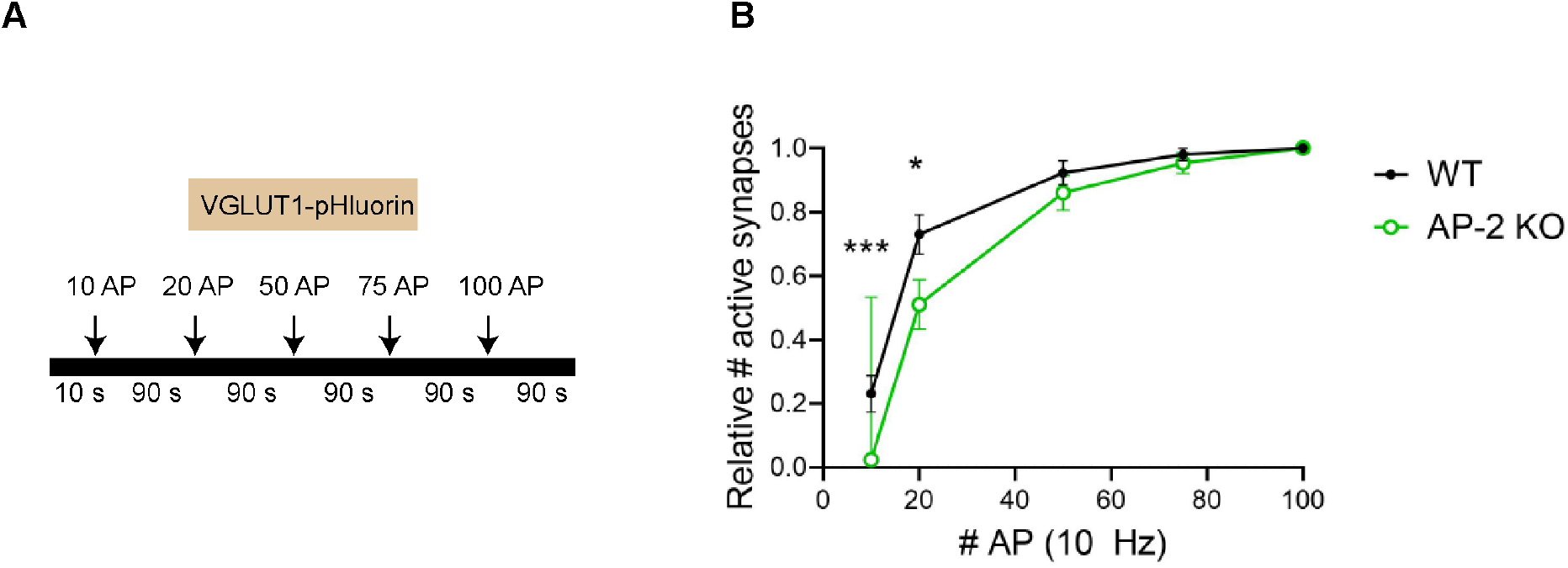
Decreased fraction of active synapses in neurons lacking AP-2. (A,B) Fraction of active synapses in AP-2 depleted neurons is reduced depending on the stimulation strength. (A) Schematic depicting the stimulation protocol of consecutive trains of AP with interstimulus intervals of 90 s by using VGLUT1-SEP. (B) Quantification of number of active synapses (signals > 4x S.D. of the noise) in WT and AP-2 KO neurons upon each AP train. Values for 100 APs were set to 1. Data represent mean ± SEM of n = 52 boutons for WT and n = 43 boutons for AP-2 KO neurons. \*\*\**p*_10AP_ = 0.0005; \**p*_20 AP_ = 0.0276; *p*_50 AP_ = 0.3275; *p*_75 AP_ = 0.4545. Two-sided unpaired *t*-test. Raw data can be found in Figure 4-figure supplement 3-source data 1.

**Figure 5–figure supplement 4.**
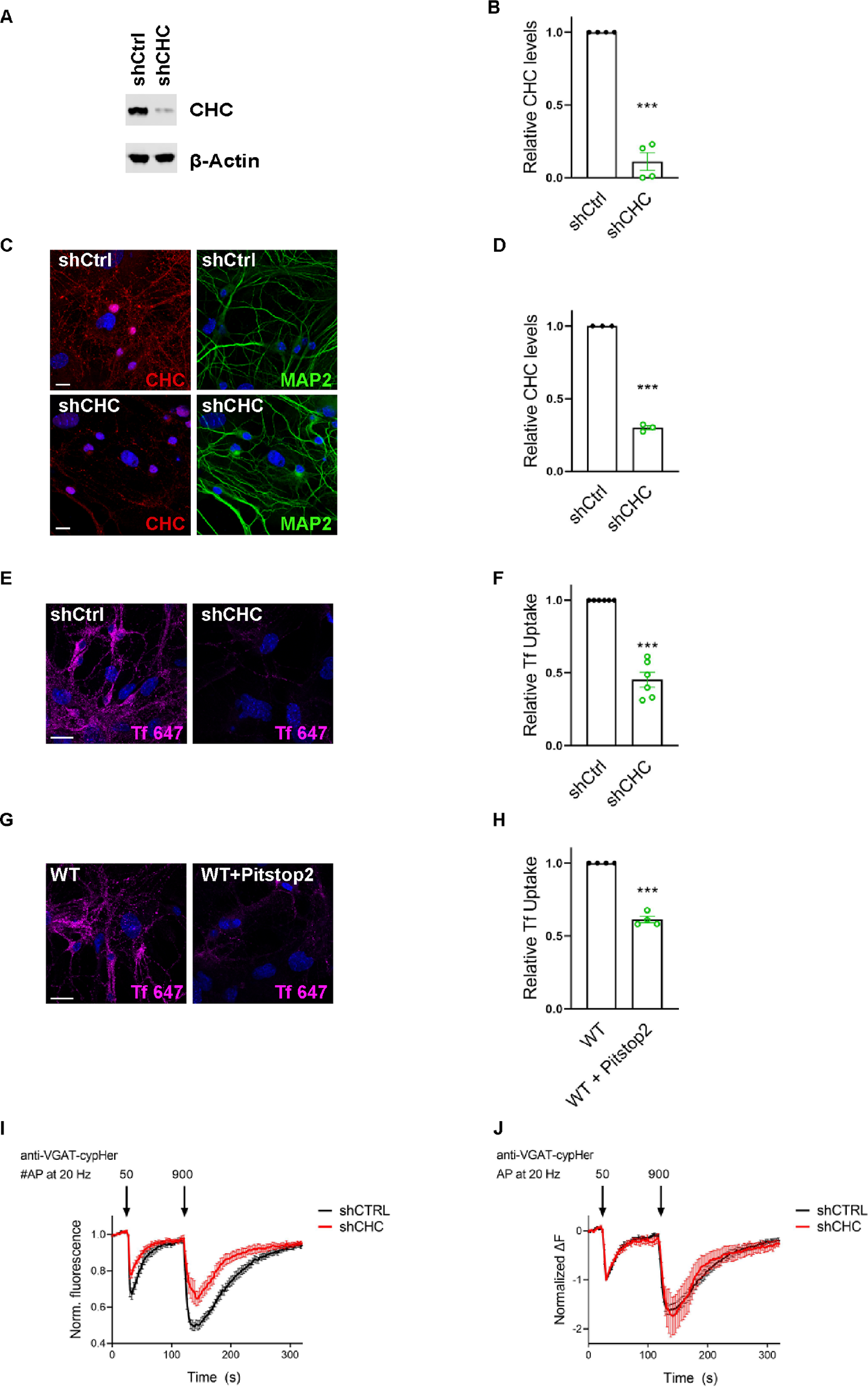
Clathrin is involved in the reformation of SVs downstream of CIE to sustain neurotransmission. (A-D) Validation of clathrin knockdown. (A) Representative immunoblot of lysates from neurons transduced with lentivirus expressing either non-specific shRNA (shCTRL) or shRNA targeting CHC (shCHC) and probed with indicated antibodies. (B) Quantification of CHC levels. Protein expression was normalized to actin. Values for shCTRL were set to 1. Data represent mean ± SEM of n = 4 independent experiments. \*\*\**p* = 0.0007. Two-sided one-sample *t*-test. (C) *Representative images* of hippocampal neuron cultures *transduced* with *lentivirus* encoding shCTRL and shCHC immunostained for CHC (red) and the neuronal marker MAP2 (green). The scale bar represents 20 μm. (D) Quantification of CHC fluorescence intensity ratio. Values for shCTRL were set to 1. Data represent mean ± SEM of n = 3 independent experiments. \*\*\**p* = 0.0004, two-sided one-sample *t*-test. (E-H) Clathrin inactivation leads to reduced CME of transferrin. (E) Representative images of primary neurons transduced with shCTRL or shCHC and allowed to internalize AlexaFluor^647^-labeled transferrin (Tf 647) for 20 min at 37°C. The scale bar represents 20 μm. (F) Quantification of data shown in (E). Values for shCTRL were set to 1. The data represent mean ± SEM from n = 6 independent experiments. \*\*\**p* = 0.0001, two-sided one-sample *t*-test. (G) Representative images of primary neurons treated either with DMSO (CTRL) or the clathrin inhibitor Pitstop 2 and allowed to internalize Tf 647 for 20 min at 37°C. Scale bar: 20 μm. (H) Quantification of data shown in (G). Values for CTRL were set to 1. The data represent mean ± SEM of n = 4 independent experiments. \*\*\**p* = 0.004, two-sided one-sample *t*-test. (I,J) Clathrin loss increases depression of neurotransmitter release without changing post-exocytic retrieval kinetics of endogenous VGAT. (I) Average normalized traces of neurons transduced with lentivirus expressing either shCTRL or shCHC, incubated with anti-VGAT cypHer-coupled antibodies and subjected to consecutive stimulus trains of 50 AP and 900 AP applied both at 20 Hz with an inter-stimulus interval of 1.5 min to determine the size of the readily releasable SV pool and the recycling SV pool. n = 10 images for shCTRL and n = 8 images for shCHC. (J) Average normalized traces of neurons transduced with lentivirus expressing either shCTRL or shCHC, incubated with anti-VGAT cypHer-coupled antibodies and subjected to consecutive stimulus trains of 50 AP and 900 AP applied both at 20 Hz with an inter-stimulus interval of 1.5 min. The fluorescence was normalized to the first peak at the end of the first AP train with 50 APs. No differences in the kinetics of endocytic recovery were observed. n = 10 images for shCTRL and n = 8 images for shCHC. Raw data can be found in Figure 5-figure supplement 4-source data 1, and Figure 5-figure supplement 4-source data 2.

**Figure 6–figure supplement 5.**
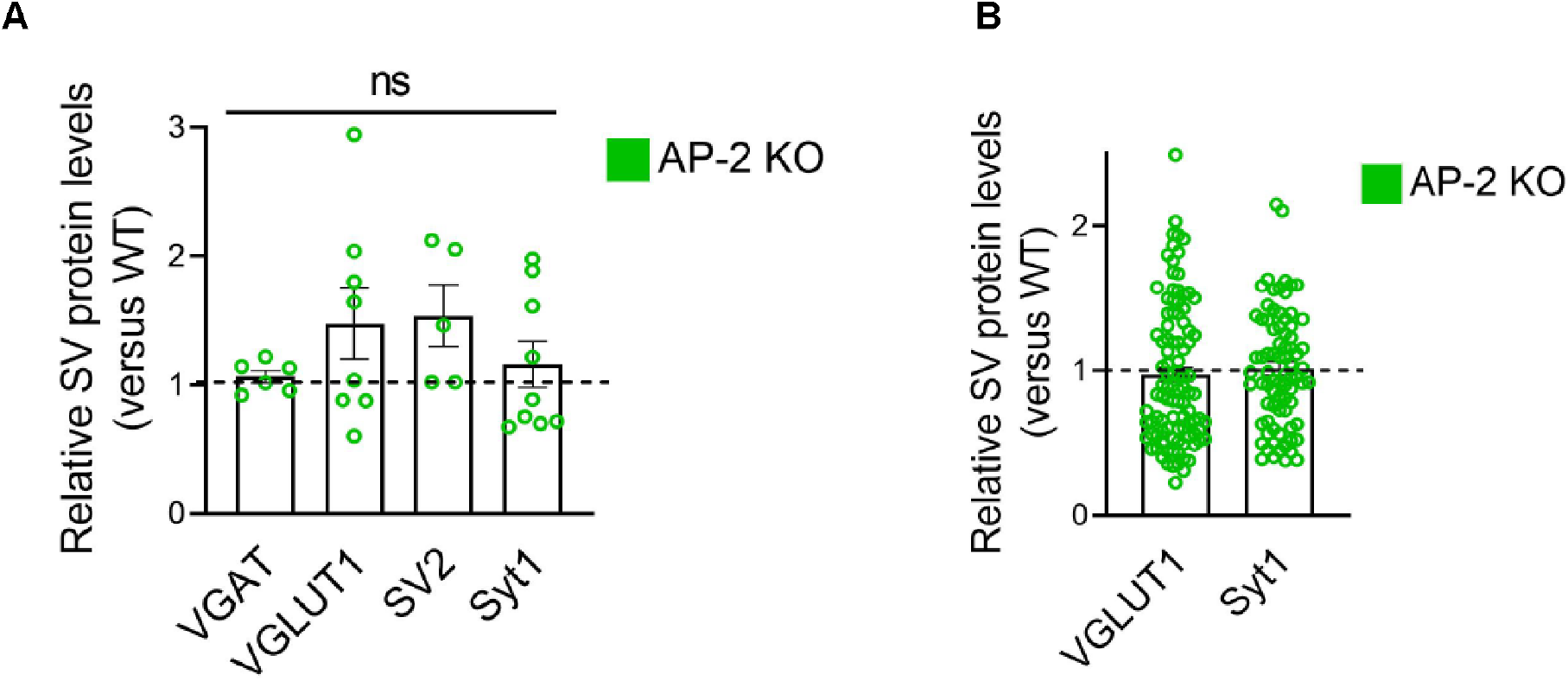
AP-2 depletion does not change the total levels of SV proteins. (A,B) Levels of presynaptic proteins are largely unaffected in the absence of AP-2. (A) Quantification of presynaptic proteins analyzed in lysates from WT and AP-2 KO neurons. Protein expression was normalized to GAPDH. Values for WT neurons were set to 1. Data represent mean ± SEM of n = 6 for VGLUT1 (*p* = 0.2383), n = 8 for VGAT (*p* = 0.1274), n = 5 for SV2 (*p* = 0.0883) and n = 9 independent experiments for Syt1 (*p* = 0.4001). Two-sided one-sample *t*-test. (B) Fluorescence intensity quantification of the presynaptic marker proteins (VGLUT1 and Syt1) analyzed from WT and AP-2 KO hippocampal neuron cultures. Values to WT. Data represent mean ± SEM of n = 99 images for VGLUT1 (*p* = 0.5547), n = 78 images for Syt1 (*p* = 0.7623). Two-sided one-sample *t*-test. Raw data can be found in Figure 6-figure supplement 5-source data 1.

**Figure 7–figure supplement 6.**
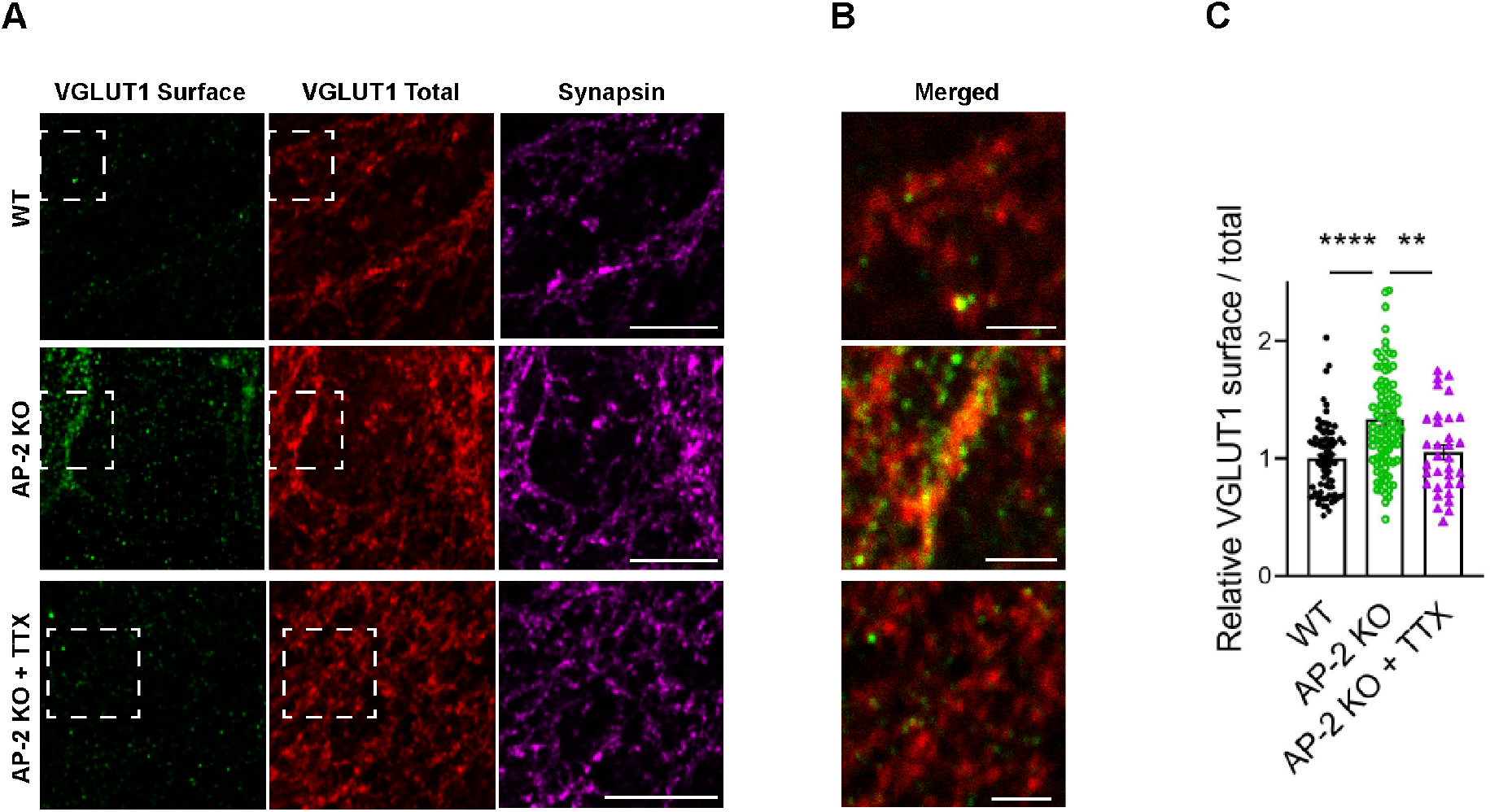
AP-2 depletion alters the localization of VGLUT1 in an activity-dependent manner. (A-C) Block of neuronal network activity rescues surface accumulation of VGLUT1 in neurons lacking AP-2. (A) Representative confocal images of WT and AP-2 KO neurons treated or not with tetrodotoxin (TTX) since DIV7 to block spontaneous action potentials and co-immunostained for total VGLUT1 (red), surface VGLUT1 (green) and the presynaptic marker synapsin (magenta). Scale bar: 10 µm. (B) A zoom of the marked areas in (A). Scale bars, 2 µm. (C) Elevated ratio of surface/total VGLUT1 in AP-2 KO neurons is rescued when neurons were treated with TTX. Values for WT neurons were set to 1. Data represent mean ± SEM of n_WT_ = 81 images, n_AP2 KO_ = 92 images and n_AP2 KO+TTX_ = 33 images. \*\*\*\**p*_WT vs AP2 KO_ < 0.0001, *p* _WT vs AP2 KO+TTX_ = 0.8004, \*\**p*_AP2 KO vs AP2 KO+TTX_ = 0.0026. One-way ANOVA with Tukey’s post-test. Raw data can be found in Figure 7-figure supplement 6-source data 1.

**Figure 8–figure supplement 7.**
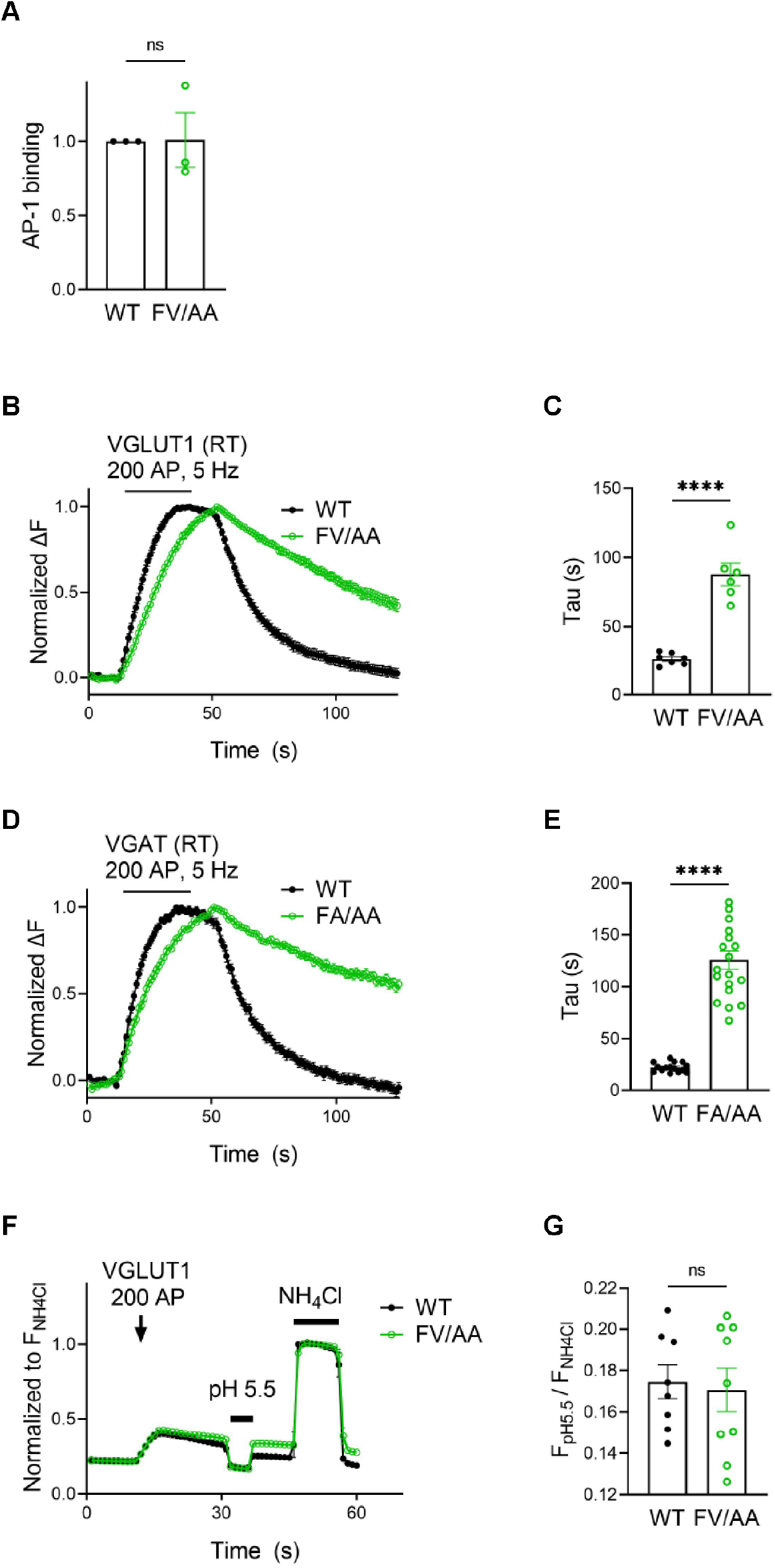
Defective retrieval of vesicular neurotransmitter transporters carrying mutations in the AP-2-binding dileucine motif cannot be ascribed to defects in acidification. (A) VGLUT1 – AP-1 interaction is insensitive to the F510A/ V511A mutation. Quantification of the band intensity corresponding to AP-1 γ subunit bound either to GST-VGLUT1-WT or VGLUT1-FV/AA mutant (from the GST-pull down assay shown in Figure 8B). Values for VGLUT1-WT were set to 1. Data represent mean ± SEM of n = 3 independent experiments. *p* = 0.9561. Two-sided one-sample *t*-test. (B-E) Endocytic defect caused by AP-2 binding deficient mutations in the vesicular transporters is aggravated under conditions that favor CME (e.g. at room temperature (RT) with low stimulation frequency). Average normalized traces of hippocampal neurons transduced with lentivirus expressing either VGLUT1-WT-SEP and VGLUT1-FV/AA-SEP (B) or VGAT-WT-SEP and VGAT-FA/AA-SEP (D) upon stimulation of 200 APs at 5 Hz at RT. Quantification of fluorescence decay (Tau) of SEP signal in VGLUT1-SEP-expressing neurons (C) (τ_VGLUT1 WT_ = 26.28 ± 1.694 s, τ_VGLUT1 FV/AA_= 87.86 ± 8.155 s) or VGAT-SEP-expressing neurons (E) (τ_VGAT WT_ = 22.45 ± 1.166 s, τ _VGAT FA/AA_= 125.6 ± 8.706 s). Data represent the mean ± SEM: VGLUT1 (n_VGLUT1 WT_ = 7 images, n_VGLUT1 FV/AA_ = 6 images; **** *p* < 0.0001) and for VGAT (n_VGAT WT_ = 15 images, n_VGAT FA/AA_ = 19 images; **** *p* < 0.0001). Two-sided unpaired *t*-test. (F,G) Slower post-stimulus retrieval of AP-2 deficient binding variant of VGLUT1 is not caused by defects in re-acidification of endocytosed vesicles. (F) Average normalized traces showing post-stimulus application (200 AP, 5 Hz) of an acid quench protocol to reveal the fraction of SV protein retrieved immediately after stimulation in hippocampal neurons transduced with lentivirus expressing either VGLUT1-WT-SEP or VGLUT1-FV/AA-SEP. Values are normalized to the corresponding maximal fluorescent peak obtained upon NH_4_Cl application. (G) Fluorescence signal ratio between minimum (acid load) and maximum (NH_4_Cl) was quantified. Data represent the mean ± SEM of n_VGLUT1 WT_ = 8 images, n_VGLUT1 FV/AA_ = 9 images. *p* = 0.7781, two-sided unpaired *t*-test. Raw data can be found in Figure 8-figure supplement 7-source data 1, and Figure 8-figure supplement 7-source data 2.

## SUPPLEMENTARY FILES

Supplementary File 1

**Supplementary Table 1:**
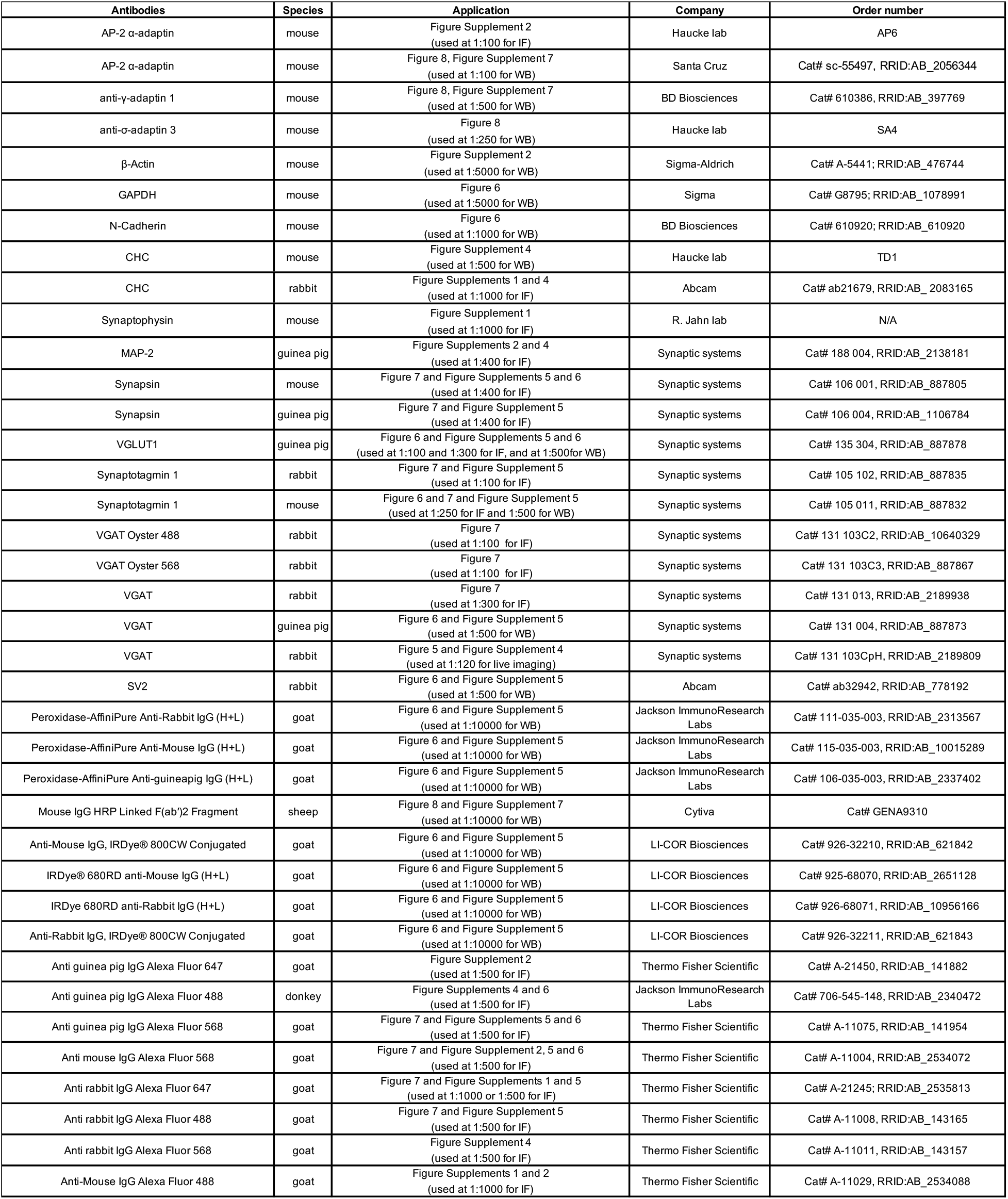
List of antibodies.

## REFERENCES

1. T. C. Sudhof, The synaptic vesicle cycle. Annu Rev Neurosci 27, 509–547 (2004).

2. J. Dittman, T. A. Ryan, Molecular circuitry of endocytosis at nerve terminals. Annu Rev Cell Dev Biol 25, 133–160 (2009).

3. N. L. Kononenko, V. Haucke, Molecular mechanisms of presynaptic membrane retrieval and synaptic vesicle reformation. Neuron 85, 484–496 (2015).

4. S. O. Rizzoli, Synaptic vesicle recycling: steps and principles. EMBO J 33, 788–822 (2014).

5. S. Takamori et al., Molecular anatomy of a trafficking organelle. Cell 127, 831–846 (2006).

6. B. G. Wilhelm et al., Composition of isolated synaptic boutons reveals the amounts of vesicle trafficking proteins. Science 344, 1023–1028 (2014).

7. I. Delvendahl, N. P. Vyleta, H. von Gersdorff, S. Hallermann, Fast, Temperature-Sensitive and Clathrin-Independent Endocytosis at Central Synapses. Neuron 90, 492–498 (2016).

8. N. L. Kononenko et al., Clathrin/AP-2 mediate synaptic vesicle reformation from endosome-like vacuoles but are not essential for membrane retrieval at central synapses. Neuron 82, 981–988 (2014).

9. T. Soykan et al., Synaptic Vesicle Endocytosis Occurs on Multiple Timescales and Is Mediated by Formin-Dependent Actin Assembly. Neuron 93, 854–866.e854 (2017).

10. S. Watanabe et al., Ultrafast endocytosis at mouse hippocampal synapses. Nature 504, 242–247 (2013).

11. S. Watanabe et al., Clathrin regenerates synaptic vesicles from endosomes. Nature 515, 228–233 (2014).

12. I. Milosevic, Revisiting the Role of Clathrin-Mediated Endoytosis in Synaptic Vesicle Recycling. Frontiers in Cellular Neuroscience 12 (2018).

13. J. E. Heuser, T. S. Reese, Evidence for recycling of synaptic vesicle membrane during transmitter release at the frog neuromuscular junction. J Cell Biol 57, 315–344 (1973).

14. T. Mitsunari et al., Clathrin adaptor AP-2 is essential for early embryonal development. Mol Cell Biol 25, 9318–9323 (2005).

15. Y. Saheki, P. De Camilli, Synaptic vesicle endocytosis. Cold Spring Harb Perspect Biol 4, a005645 (2012).

16. M. Kaksonen, A. Roux, Mechanisms of clathrin-mediated endocytosis. Nat Rev Mol Cell Biol 19, 313–326 (2018).

17. B. Granseth, B. Odermatt, S. J. Royle, L. Lagnado, Clathrin-mediated endocytosis is the dominant mechanism of vesicle retrieval at hippocampal synapses. Neuron 51, 773–786 (2006).

18. B. Granseth, B. Odermatt, S. J. Royle, L. Lagnado, Clathrin-mediated endocytosis: the physiological mechanism of vesicle retrieval at hippocampal synapses. J Physiol 585, 681–686 (2007).

19. E. Alés et al., High calcium concentrations shift the mode of exocytosis to the kiss-and- run mechanism. Nature Cell Biology 1, 40–44 (1999).

20. W. Shin et al., Visualization of Membrane Pore in Live Cells Reveals a Dynamic-Pore Theory Governing Fusion and Endocytosis. Cell 173, 934–945 e912 (2018).

21. J. Kasprowicz, S. Kuenen, J. Swerts, K. Miskiewicz, P. Verstreken, Dynamin photoinactivation blocks Clathrin and alpha-adaptin recruitment and induces bulk membrane retrieval. J Cell Biol 204, 1141–1156 (2014).

22. J. Kasprowicz et al., Inactivation of clathrin heavy chain inhibits synaptic recycling but allows bulk membrane uptake. J Cell Biol 182, 1007–1016 (2008).

23. H. Heerssen, R. D. Fetter, G. W. Davis, Clathrin dependence of synaptic-vesicle formation at the Drosophila neuromuscular junction. Curr Biol 18, 401–409 (2008).

24. C. Imig et al., Ultrastructural Imaging of Activity-Dependent Synaptic Membrane-Trafficking Events in Cultured Brain Slices. Neuron 108, 843–860 e848 (2020).

25. S. H. Kim, T. A. Ryan, Synaptic vesicle recycling at CNS synapses without AP-2. J Neurosci 29, 3865–3874 (2009).

26. S. M. Voglmaier et al., Distinct endocytic pathways control the rate and extent of synaptic vesicle protein recycling. Neuron 51, 71–84 (2006).

27. M. S. Santos, C. K. Park, S. M. Foss, H. Li, S. M. Voglmaier, Sorting of the vesicular GABA transporter to functional vesicle pools by an atypical dileucine-like motif. J Neurosci 33, 10634–10646 (2013).

28. H. Li, M. S. Santos, C. K. Park, Y. Dobry, S. M. Voglmaier, VGLUT2 Trafficking Is Differentially Regulated by Adaptor Proteins AP-1 and AP-3. Front Cell Neurosci 11, 324 (2017).

29. S. M. Foss, H. Li, M. S. Santos, R. H. Edwards, S. M. Voglmaier, Multiple dileucine-like motifs direct VGLUT1 trafficking. J Neurosci 33, 10647–10660 (2013).

30. M. A. Cousin, Integration of Synaptic Vesicle Cargo Retrieval with Endocytosis at Central Nerve Terminals. Frontiers in cellular neuroscience 11, 234–234 (2017).

31. N. Kaempf et al., Overlapping functions of stonin 2 and SV2 in sorting of the calcium sensor synaptotagmin 1 to synaptic vesicles. Proceedings of the National Academy of Sciences 112, 7297 (2015).

32. S. J. Koo et al., Vesicular Synaptobrevin/VAMP2 Levels Guarded by AP180 Control Efficient Neurotransmission. Neuron 88, 330–344 (2015).

33. Y. Mori, S. Takamori, Molecular Signatures Underlying Synaptic Vesicle Cargo Retrieval. Front Cell Neurosci 11, 422 (2017).

34. K. Takei, O. Mundigl, L. Daniell, P. De Camilli, The synaptic vesicle cycle: a single vesicle budding step involving clathrin and dynamin. J Cell Biol 133, 1237–1250 (1996).

35. G. Miesenbock, D. A. De Angelis, J. E. Rothman, Visualizing secretion and synaptic transmission with pH-sensitive green fluorescent proteins. Nature 394, 192–195 (1998).

36. S. Sankaranarayanan, D. De Angelis, J. E. Rothman, T. A. Ryan, The use of pHluorins for optical measurements of presynaptic activity. Biophys J 79, 2199–2208 (2000).

37. P. P. Atluri, T. A. Ryan, The kinetics of synaptic vesicle reacidification at hippocampal nerve terminals. J Neurosci 26, 2313–2320 (2006).

38. Y. Egashira, M. Takase, S. Takamori, Monitoring of vacuolar-type H+ ATPase-mediated proton influx into synaptic vesicles. J Neurosci 35, 3701–3710 (2015).

39. L. von Kleist et al., Role of the clathrin terminal domain in regulating coated pit dynamics revealed by small molecule inhibition. Cell 146, 471–484 (2011).

40. Y. Hua et al., A readily retrievable pool of synaptic vesicles. Nat Neurosci 14, 833–839 (2011).

41. V. N. Murthy, C. F. Stevens, Reversal of synaptic vesicle docking at central synapses. Nat Neurosci 2, 503–507 (1999).

42. F. Opazo et al., Limited intermixing of synaptic vesicle components upon vesicle recycling. Traffic 11, 800–812 (2010).

43. J. S. Bonifacino, L. M. Traub, Signals for Sorting of Transmembrane Proteins to Endosomes and Lysosomes. Annual Review of Biochemistry 72, 395–447 (2003).

44. B. T. Kelly et al., A structural explanation for the binding of endocytic dileucine motifs by the AP2 complex. Nature 456, 976–979 (2008).

45. I. Helbig et al., A Recurrent Missense Variant in AP2M1 Impairs Clathrin-Mediated Endocytosis and Causes Developmental and Epileptic Encephalopathy. Am J Hum Genet 104, 1060–1072 (2019).

46. H. T. McMahon, E. Boucrot, Molecular mechanism and physiological functions of clathrin-mediated endocytosis. Nat Rev Mol Cell Biol 12, 517–533 (2011).

47. Z. Taoufiq et al., Hidden proteome of synaptic vesicles in the mammalian brain. Proc Natl Acad Sci U S A 117, 33586–33596 (2020).

48. M. Orlando, D. Schmitz, C. Rosenmund, M. A. Herman, Calcium-Independent Exo-endocytosis Coupling at Small Central Synapses. Cell Rep 29, 3767–3774 e3763 (2019).

49. N. L. Kononenko et al., Compromised fidelity of endocytic synaptic vesicle protein sorting in the absence of stonin 2. Proc Natl Acad Sci U S A 110, E526–535 (2013).

50. S. E. Lee, S. Jeong, U. Lee, S. Chang, SGIP1alpha functions as a selective endocytic adaptor for the internalization of synaptotagmin 1 at synapses. Mol Brain 12, 41 (2019).

51. D. T. Stimson et al., Drosophila stoned proteins regulate the rate and fidelity of synaptic vesicle internalization. J Neurosci 21, 3034–3044 (2001).

52. A. M. Phillips, M. Ramaswami, L. E. Kelly, Stoned. Traffic 11, 16–24 (2010).

53. N. Jung et al., Molecular basis of synaptic vesicle cargo recognition by the endocytic sorting adaptor stonin 2. J Cell Biol 179, 1497–1510 (2007).

54. P. Y. Pan, J. Marrs, T. A. Ryan, Vesicular glutamate transporter 1 orchestrates recruitment of other synaptic vesicle cargo proteins during synaptic vesicle recycling. J Biol Chem 290, 22593–22601 (2015).

55. K. I. Willig, S. O. Rizzoli, V. Westphal, R. Jahn, S. W. Hell, STED microscopy reveals that synaptotagmin remains clustered after synaptic vesicle exocytosis. Nature 440, 935–939 (2006).

56. D. R. Lazzell, R. Belizaire, P. Thakur, D. M. Sherry, R. Janz, SV2B regulates synaptotagmin 1 by direct interaction. J Biol Chem 279, 52124–52131 (2004).

57. A. E. Schivell, R. H. Batchelor, S. M. Bajjalieh, Isoform-specific, calcium-regulated interaction of the synaptic vesicle proteins SV2 and synaptotagmin. J Biol Chem 271, 27770–27775 (1996).

58. J. Yao, A. Nowack, P. Kensel-Hammes, R. G. Gardner, S. M. Bajjalieh, Cotrafficking of SV2 and synaptotagmin at the synapse. J Neurosci 30, 5569–5578 (2010).

59. S. A. Mutch et al., Protein quantification at the single vesicle level reveals that a subset of synaptic vesicle proteins are trafficked with high precision. J Neurosci 31, 1461–1470 (2011).

60. O. Martzoukou, S. Amillis, A. Zervakou, S. Christoforidis, G. Diallinas, The AP-2 complex has a specialized clathrin-independent role in apical endocytosis and polar growth in fungi. Elife 6 (2017).

61. R. Pascolutti et al., Molecularly Distinct Clathrin-Coated Pits Differentially Impact EGFR Fate and Signaling. Cell Rep 27, 3049–3061 e3046 (2019).

62. Y. Egashira et al., Unique pH dynamics in GABAergic synaptic vesicles illuminates the mechanism and kinetics of GABA loading. Proc Natl Acad Sci U S A 113, 10702–10707 (2016).

63. M. K. Diril, M. Wienisch, N. Jung, J. Klingauf, V. Haucke, Stonin 2 is an AP-2-dependent endocytic sorting adaptor for synaptotagmin internalization and recycling. Dev Cell 10, 233–244 (2006).

64. S. E. Kwon, E. R. Chapman, Synaptophysin regulates the kinetics of synaptic vesicle endocytosis in central neurons. Neuron 70, 847–854 (2011).

65. S. Y. Kawaguchi, T. Sakaba, Control of inhibitory synaptic outputs by low excitability of axon terminals revealed by direct recording. Neuron 85, 1273–1288 (2015).

66. H. Hioki et al., High-level transgene expression in neurons by lentivirus with Tet-Off system. Neurosci Res 63, 149–154 (2009).

67. D. Cai, V. M. Latham, Jr., X. Zhang, G. I. Shapiro, Combined depletion of cell cycle and transcriptional cyclin-dependent kinase activities induces apoptosis in cancer cells. Cancer Res 66, 9270–9280 (2006).

68. M. Wienisch, J. Klingauf, Vesicular proteins exocytosed and subsequently retrieved by compensatory endocytosis are nonidentical. Nat Neurosci 9, 1019–1027 (2006).

69. S. E. Kwon, E. R. Chapman, Glycosylation is dispensable for sorting of synaptotagmin 1 but is critical for targeting of SV2 and synaptophysin to recycling synaptic vesicles. J Biol Chem 287, 35658–35668 (2012).

70. T. Lopez-Hernandez, D. Puchkov, E. Krause, T. Maritzen, V. Haucke, Endocytic regulation of cellular ion homeostasis controls lysosome biogenesis. Nat Cell Biol 22, 815–827 (2020).

71. M. Jiang, G. Chen, High Ca2+-phosphate transfection efficiency in low-density neuronal cultures. Nat Protoc 1, 695–700 (2006).

72. J. Schindelin et al., Fiji: an open-source platform for biological-image analysis. Nat Methods 9, 676–682 (2012).

73. P. Virtanen et al., SciPy 1.0: fundamental algorithms for scientific computing in Python. Nat Methods 17, 261–272 (2020).

